# Maternal sterol 27-hydroxylase is crucial for securing fetal development

**DOI:** 10.1101/2023.11.08.566330

**Authors:** Mitsuyoshi Suzuki, Satoshi Nakano, Natsumi Miharada, Hajime Takei, Pavan Prabhala, Mark van der Garde, Catharina Müller, Valgardur Sigurdsson, Maolake Aerken, Kiyoka Saito, Shuhei Koide, Gunilla Westergren-Thorsson, Mattias Magnusson, Genta Kakiyama, Hiroshi Nittono, Kenichi Miharada

**Affiliations:** Division of Molecular Medicine and Gene Therapy, Lund Stem Cell Center, Lund University, 221 48 Lund, Sweden; Department of Pediatrics, Juntendo University Faculty of Medicine, 113-8421 Tokyo, Japan; International Research Center for Medical Sciences, Kumamoto University, 860-0811 Kumamoto, Japan; Junshin Clinic Bile Acid Institute, 152-0011 Tokyo, Japan; Department of Medicine III, Hematology and Oncology, Technical University of Munich, D-81675 Munich, Germany; Department of Experimental Medical Science, Lund University, 221 48 Lund, Sweden; Division of Stem Cell and Molecular Medicine, Center for Stem Cell Biology and Regenerative Medicine, The Institute of Medical Science, The University of Tokyo, 108-8639 Tokyo, Japan; Department of Internal Medicine, Virginia Commonwealth University School of Medicine, Richmond, 23298 VA; Research Services, Central Virginia Veterans Affairs Healthcare System, Richmond, 23298 VA

**Author notes:** Correspondence Kenichi Miharada.

## Abstract

The maternal body helps in providing nutrients and degrading toxic metabolites instead of the fetal body; disruptions in these mechanisms affect normal fetal development. Sterol 27-hydroxylase (*Cyp27a1*) is involved in the alternative pathway of bile acid synthesis, which is enhanced during pregnancy. However, its role in fetal development remains unclear. Here, we demonstrate that maternal Cyp27a1 activity is essential for progression of normal pregnancy and fetal organ formation. Depletion of maternal *Cyp27a1* reduced the pregnancy rate and litter size. Newborn mice died of respiratory distress syndrome resulting from the absence of mature alveolar epithelial cells. These phenotypes were caused by 7α-hydroxycholesterol (7α-HC) accumulating in *Cyp27a1*-deficient mice. Mechanistically, 7α-HC destabilized the Fau protein, mediating ribosome assembly, the downregulation of which caused poor polysome formation, lower protein synthesis, and impaired lung maturation. Overall, this study revealed an essential mechanism of securing fetal development by degrading a toxic metabolite in the maternal body.

## Introduction

During pregnancy, the maternal circulation plays a vital role in providing essential factors and degrading toxic metabolites instead of developing fetuses. The disruptions of these mechanisms may critically affect normal fetal growth and development. Metabolites transmitted from the maternal body to the embryonal/fetal body through the placenta can be both crucial regulators and critical insults (Bowman et al., 2021; Villar et al., 2022; Gleason et al., 2023). The fetal livers are not fully matured, requiring the maternal body to compensate for essential metabolic activity during the pregnancy. For instance, the degradation of aldehydes naturally produced in the developing fetuses by the maternal body helps in fetal brain development and protects bone marrow cells from DNA damage (Langevin et al., 2011; Oberbeck et al., 2014). Furthermore, lowering iron-dependent oxidative stress levels is imperative for reducing the risks of preterm birth (Sakata et al., 2008). Thus, managing metabolic conditions is crucial for an effective fetal-maternal crosstalk.

Cholesterol is the structural component of cell membranes and is a precursor for vitamin D and several steroid hormones (Woollett, 2011; Ellinger and Chatuphonprasert, 2022). Bile acids (BAs) are the major hepatic cholesterol metabolites that are synthesized via oxysterols through multiple enzymatic reactions. Even though BAs are primarily involved in lipid absorption, they also function as molecular chaperones and signaling molecules (Özcan et al., 2006; Watanabe et al., 2006; Nguyen and Bouscarel, 2008; de Aguiar Vallim et al., 2013). However, dysregulated BAs and oxysterols could pose toxic insults (Pataia et al., 2017; Song et al., 2021).

BA synthesis involves two main synthetic pathways: the classic pathway (neutral pathway) initiated by cholesterol 7α-hydroxylase (Cyp7a1) and the alternative pathway (acidic pathway) initiated by sterol 27-hydroxylase (Cyp27a1) (Hofmann, 1999). Cyp7a1 is specifically expressed in the liver and generates 7α-hydroxycholesterol (7α-HC) for pushing the cholesterol catabolism to BAs. In contrast, Cyp27a1 is broadly expressed in many organs and cells throughout the body and exerts multiple enzymatic reactions, including vitamin D metabolism. Cyp27a1 is involved in the hydroxylation of multiple intermediary sterols and BAs, both in the classic pathway and the alternative pathway (Figure 1A and Supplementary Figure S1A) (Griffiths et al., 2019). In the initial step of the alternative pathway, Cyp27a1 primarily catalyzes 27-hydroxylation of cholesterol to generate 27-hydroxycholesterol (27-HC), which can be further oxidized to 3β-hydroxy-5-cholestenoic acid (3βHCA) (Lorbek et al., 2011). Note that the systemic name of the Cyp27a1-metabolite is (25R)26-hydroxycholesterol (Fakheri and Javitt, 2012); however, we used the historical name of 27-HC in this paper.

**Figure 1.**
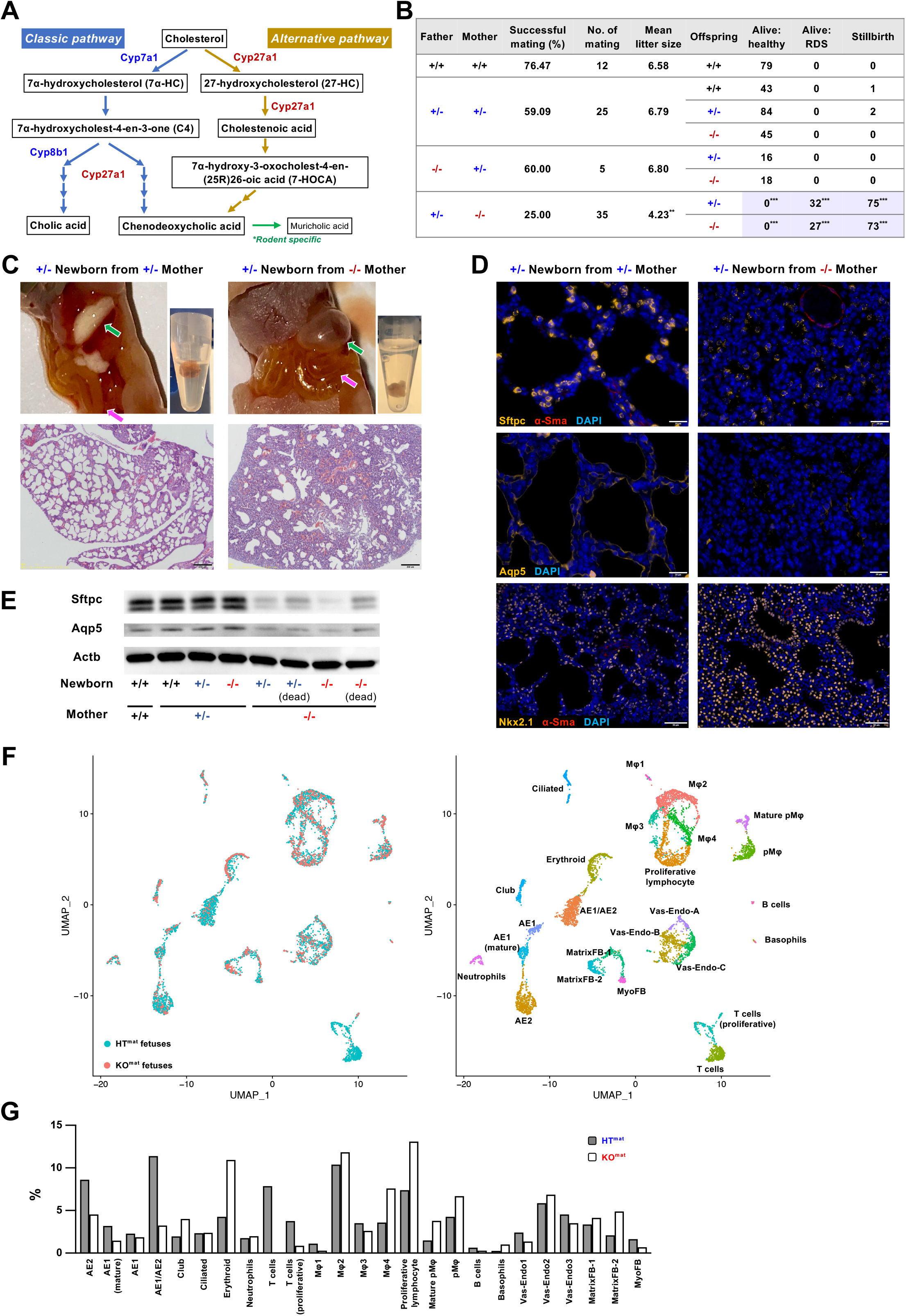
Newborns delivered from *Cyp27a1^-/-^* mothers die due to RDS. **(A)** Simplified BA synthetic pathways. The classic pathway is shown in blue arrows, and the alternative pathway is shown in yellow arrows. The rodent specific muricholic acid pathway is shown in a green arrow. **(B)** Outcomes from the mating of *Cyp27a1* deficient mice. WT (+/+), HT (+/-), or KO (-/-) knockout mice were crossed in various combinations. The successful mating rate was calculated by the number of successful pregnancies divided by the number of timed mating attempts. Litter size was the sum of live births and stillbirths from one pregnant mouse. Respiratory distress syndrome was determined by the observation of gasping respiration. **(C)** Representative photos of newborn mice delivered from *Cyp27a1^+/-^*and *Cyp27a1^-/-^* mothers. The top left panels: the abdomen of the newborn mice. The green arrows indicate the stomach, and the pink arrows indicate the intestine. The top right panels: the lungs of the newborn mice put into PBS. The bottom panels: H&E staining of the tissue sections of the newborn mice’s lungs. The scale bar represents 200 μm. **(D)** Immunostaining of the tissue sections of the newborn mice’s lungs. The top panels: staining of Sftpc (yellow), α-Sma (red), and DAPI (blue). The scale bar represents 20 μm. The middle panels: Aqp5 (yellow) and DAPI (blue). The scale bar represents 20 μm. The bottom panels: Nkx2.1 (yellow), α-Sma (red), and DAPI (blue). The scale bar represents 50 μm. **(E)** Western blot analyses of Sftpc, Aqp5, and Actb in the lung tissues of fetuses delivered from different mothers. **(F)** scRNA-seq analysis of lung tissues from E18.5 *Cyp27a^-^* HT^mat^ and KO^mat^ KO fetuses. UMAP plots of the cells from three embryos are shown. AE1: type I alveolar epithelial cells, mature AE1: mature type I alveolar epithelial cells, AE2: type II alveolar epithelial cells, AE1/AE2: type I & II common alveolar epithelial progenitors, ciliated: ciliated cells, club: club cells, Vas-Endo-1: vascular endothelial cells 1, Vas-Endo-2: vascular endothelial cells 2, Vas-Endo-3: vascular endothelial cells 3, MatrixFB-1: matrix fibroblast 1, MatrixFB-2: matrix fibroblast 2, MyoFB: myofibroblast, Mφ1: macrophage-1, Mφ2: macrophage-2, Mφ3: macrophage-3, Mφ4: macrophage-4, pMφ: primitive macrophages, mature pMφ: mature primitive macrophages, erythroid: erythroid cells, proliferative lymphocyte, neutrophils, T cells, proliferative T cells, B cells, basophils. **(G)** Frequencies of each cluster of *Cyp27a^-^* HT^mat^ and KO^mat^ KO fetuses. \*\**p* < 0.01, \*\*\**p* < 0.001.

The alternative pathway is reported to be more active during embryogenesis (Nakagawa and Setchell, 1990; Itoh and Onishi, 2000; Marin et al., 2008; Sato et al., 2020). In humans, the plasma of pregnant women shows increased 27-HC concentration during the first trimester (Winkler et al., 2017); however, the physiological significance of enhanced alternative pathway activity, especially that of Cyp27a1, remains largely unidentified. We have previously demonstrated that maternal depletion of Cyp27a1 in mice decreased fetal BA, leading to failure in the expansion of hematopoietic stem cells (HSCs) in the fetal liver due to the induction of endoplasmic reticulum (ER) stress (Sigurdsson et al., 2016), indicating the significance of maternal BA metabolism in embryonal/fetal development. Notably, it has been reported that depletion of Cyp27a1 causes severe accumulation of specific oxysterols, such as 7α-HC and 7α-Hydroxycholest-4-en-3-one (C4), and decreased 27-HC and BA levels (Griffiths et al., 2019). Oxysterols, which are oxygenated cholesterol derivatives, are intermediate products of BA synthesis and play crucial roles in lipid metabolism, immune responses, and regulation of the central nervous systems (Brown and Jessup, 1999; Schroepfer Jr. 2000; Hannedouche et al., 2011; Liu et al., 2011; Theofilopoulos et al., 2013; Dang et al., 2017; Theofilopoulos et al., 2019). Although their implications determining the fate of stem/progenitor cells in embryonic development have also been reported (Nachtergaele et al., 2012; Raleigh et al., 2018), the influences and functions of oxysterols in fetal development are still unclear.

Therefore, in this study, we aimed to analyze the role of Cyp27a1 and maternal BA synthesis in fetal development and the associated molecular mechanisms.

## Results

### Newborns delivered from *Cyp27a1^-/-^* mothers die of respiratory distress syndrome

To elucidate the effect of maternal *Cyp27a1* depletion on late fetal development, we analyzed newborns delivered from *Cyp27a1-*deficient mothers. As expected, breeding heterozygous knockout (HT, *Cyp27a1^+/-^*) mice produced wild-type (WT, *Cyp27a1^+/+^*), *Cyp27a1^+/-^*, and homozygous knockout (KO, *Cyp27a1^-/-^*) mice in Mendelian ratios (Figure 1B). *Cyp27a1^-/-^* newborns did not develop severe abnormalities when fed a regular diet and grew until reproductive age as previously reported (Rosen et al., 1998; Sigurdsson et al., 2016). However, mating *Cyp27a1^-/-^* female mice and *Cyp27a1^+/-^* male mice was less efficient, as only 25 % of the matings successfully led to birth to newborns. In comparison, the success rate of the other mating combinations was approximately 60 % or higher. The number of litters from *Cyp27a1^-/-^*mothers was significantly lower than that from *Cyp27a1^+/-^* mating (4.23 vs 6.79, respectively), and 71.5 % of the newborns were delivered dead (Figure 1B). No significant differences were observed in the offspring sex ratios (Supplementary Figure S1B). Furthermore, live newborns presented with symptoms of respiratory distress syndrome (RDS) (Figure 1B and Supplementary Movies 1 and 2) and died within 24 hours. Upon dissection, milk was confirmed in the stomachs of newborns from *Cyp27a1^+/-^* mothers (HT^mat^) half a day after birth, and their lungs floated in PBS. In contrast, no milk was observed in the newborns from *Cyp27a1^-/-^* mother (KO^mat^); instead, the stomachs were filled with air (Figure 1C). None of the newborn KO^mat^ lungs floated in PBS, and few newborns showed intestinal distension with air, suggesting respiratory system abnormalities.

Next, we histologically analyzed lung tissue sections with hematoxylin and eosin (H&E) staining and observed an underdeveloped alveolar structure in the KO^mat^ (Figures 1C and Supplementary Figure S1C). Importantly, KO^mat^ exhibited similar abnormalities regardless of genotype. However, *Cyp27a1^-/-^* newborns delivered by *Cyp27a1^+/-^* mothers, even when they were crossed with *Cyp27a1^-/-^* male mice, showed no abnormalities (Figure 1B and Supplementary Figure S1C), indicating the significance of maternal depletion of *Cyp27a1* depletion in triggering low pregnancy, small litter size, and lung abnormalities. These observations were further confirmed by *in vitro* fertilization (IVF). Fertilized eggs prepared using the sperm and eggs from *Cyp27a1^-/-^* males and *Cyp27a1^-/-^* females were transplanted into the uteri of surrogate mothers. Eighty-four mice were born with no observable RDS (Supplementary Movie 3), indicating that the lung abnormalities could be attributed to the maternal microenvironment.

### *Cyp27a1* KO^mat^ fetuses lack functional AE cells

Next, we immunostained the lung tissues from KO^mat^ newborns and observed a significant decrease in the number of Sftpc^+^ type II alveolar epithelial (AE2) cells that secrete surfactants and Aqp5^+^ type I alveolar epithelial (AE1) cells that are responsible for gas exchange (Frank et al., 2019) (Figure 1D). Conversely, the number of cells expressing the marker for precursor cells, Nkx2.1 (Ikonomou et al., 2020), showed no significant difference (Figure 1D), indicating the absence of mature AE cells. We further confirmed the loss of functional AE cells via western blot analysis. The assay showed that the levels of Sftpc and Aqp5 proteins were lower in KO^mat^ newborns than in WT^mat^ and HT^mat^ newborns (Figure 1E and Supplementary Figure S1D). To clarify whether the reduced levels of mature marker proteins were due to insufficient expression offunctional genes or the absence of mature AE cells, we performed single-cell RNA sequencing analysis (scRNA-seq) on the lung tissues from E18.5 HT^mat^ and KO^mat^ fetuses. scRNA-seq successfully identified previously reported cell populations (Guo et al., 2019; Li et al., 2020), including peripheral and proximal epithelial cells (AE1, mature AE1, AE2, AE1/AE2, ciliated cells, and club cells), vascular endothelial cells (Vas-Endo-1, -2, -3), matrix fibroblasts (Matrix-FB-1, -2), myofibroblasts (MyoFB), as well as various hematopoietic cells (macrophages (Mφ) 1-4, primitive macrophages (pMφ), mature pMφ, proliferative lymphocytes, erythroid cells, neutrophils, T cells, proliferative T cells, B cells, basophils) (Figure 1F and Supplementary Figure S2 and Supplementary Table 1). Moreover, in KO^mat^ fetuses, both AE1 and AE2 cells and their progenitor population (AE1/AE2) were vastly decreased compared to HT^mat^ fetuses (Figures 1F and G and Supplementary Figure S2). In contrast, the number of airway epithelial cells (club and ciliated cells) did not change significantly, suggesting that the reduction in the number of AE cells was a specific event. T cells numbers were also largely decreased, whereas other blood cells were unaffected (Figures 1F and G).

### Cell cycle progression is impaired in lung EpCAM^+^ cells of *Cyp27a1* KO^mat^ fetuses

To determine precisely when the critical defect in KO^mat^ newborns is initiated, we analyzed the embryos at different time points. At E14.5, all *Cyp27a1* KO^mat^ fetuses appeared identical to the WT and *Cyp27a1* HT^mat^ fetuses (Figure 2A). However, a few E16.5 *Cyp27a1* KO^mat^ fetuses exhibited bleeding from the deep neck, and subcutaneous hemorrhage spread diffusely over the body, suggesting the occurrence of the critical events during E14.5-16.5 (Figure 2A). We also noticed that many E16.5 *Cyp27a1* KO^mat^ hydropathic fetuses had fetal nuchal edema in their subcutaneous back (Supplementary Figure S3), which is associated with various fetal abnormalities, including defective lymphatic vascular development, cardiac anomalies, and anemia (Sugiyama and Hirashima, 2022). The bleeding phenotype was more evident in the E18.5 embryos than in the E16.5 embryos. However, half of the *Cyp27a1* KO^mat^ fetuses still appeared normal, even though all newborns died of RDS after birth, suggesting the occurrence of multiple abnormalities in these fetuses.

**Figure 2.**
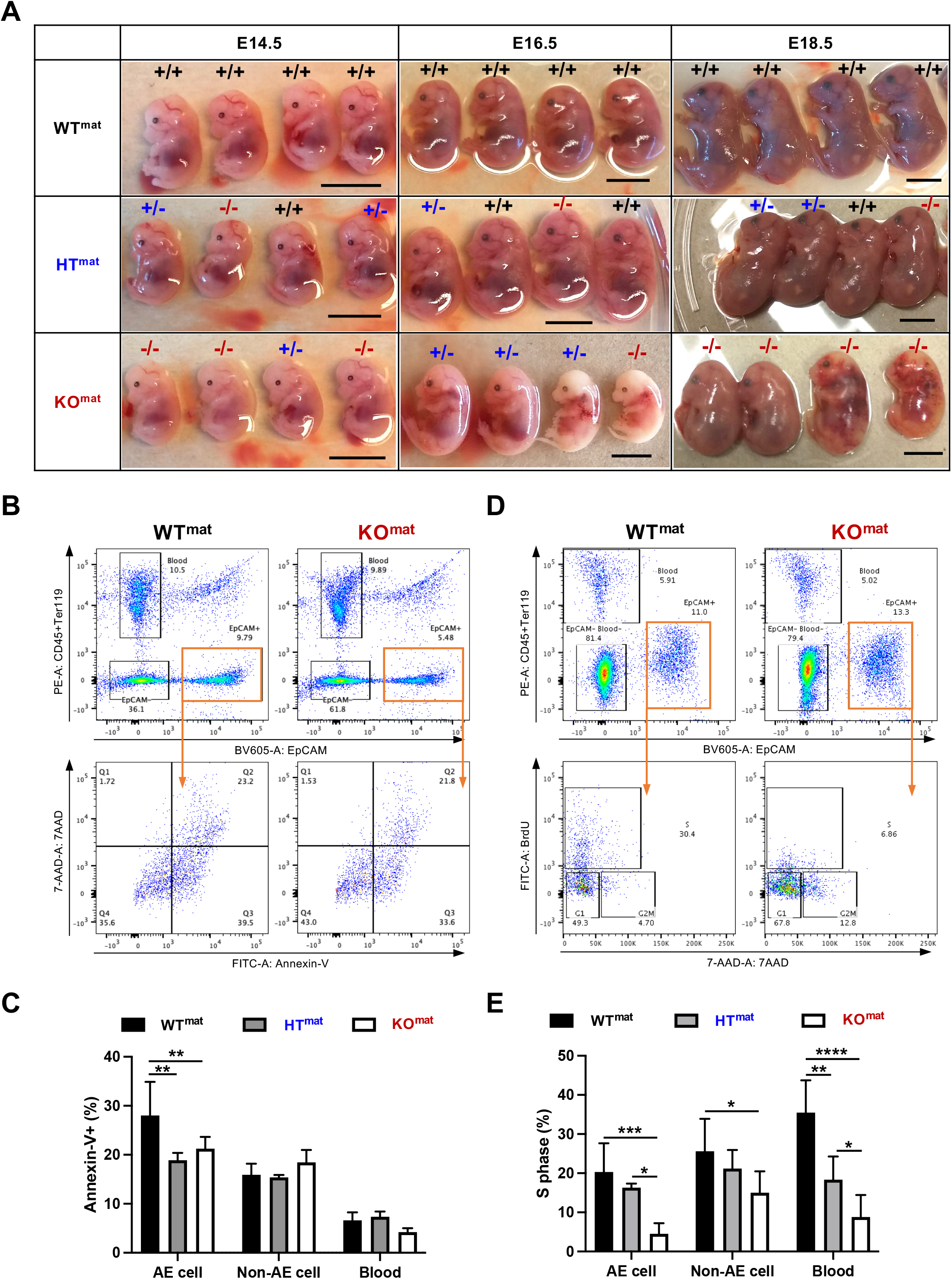
Impaired cell cycle progression in lung EpCAM^+^ cells of *Cyp27a1* KO^mat^ fetuses. **(A)** Representative images of embryos at E14.5, E16.5, and E18.5. The scale bar represents 1 cm. **(B)** A representative flow cytometry analysis plot of Annexin-V staining in the fetal lung cells. Cells were separated into CD45^+^EpCAM^-^ cells (blood), CD45^-^EpCAM^-^ cells (non-epithelial), and CD45^-^ EpCAM^+^ cells (epithelial). **(C)** Frequencies of Annexin-V^+^ cells in each fraction. **(D)** A representative flow cytometry analysis plot of BrdU staining in the fetal lung cells. **(E)** Frequencies of BrdU^+^ cells in each fraction. \**p* < 0.05, \*\**p* < 0.01, \*\*\**p* < 0.001, \*\*\*\**p* < 0.0001.

Next, we investigated whether the absence of AE cells resulted from either enhanced apoptosis or cell cycle cessation. AE cells were identified by CD45 negativity and epithelial cell adhesion molecule (EpCAM) positivity (CD45^-^EpCAM^+^ cells) (Figure 2B). The flow cytometry analysis separated fetal lung cells into CD45^+^EpCAM^-^ cells, CD45^-^EpCAM^-^ cells, and CD45^-^EpCAM^+^ cells. No significant differences in the frequency of these three populations were observed between *Cyp27a1* WT^mat^, HT^mat^, and KO^mat^ fetuses. The Annexin-V assay showed no significant increase in the number of apoptotic cells in the CD45^-^EpCAM^+^ cell population of *Cyp27a1* KO^mat^ fetuses (Figures 2B and C). In contrast, the BrdU incorporation assay revealed that the number of EpCAM^+^ cells undergoing cell cycle progression was significantly reduced in the lungs (Figures 2D and E), indicating that functional alveolar epithelial cells were deficient owing to the termination of alveolar (precursor) cell proliferation.

### Low levels of polysome formation are seen in lung cells of *Cyp27a1* KO^mat^ fetuses

To discover the mechanism underlying the disruption in the middle phase of fetal development in *Cyp27a1* KO^mat^ fetuses, we performed scRNA-seq on lung cells of E14.5 HT^mat^ and KO^mat^ fetuses. We identified 22 cell types based on a previous report (Guo et al., 2019) and the analysis using ToppFun (https://toppgene.cchmc.org/enrichment.jsp) (Figure 3A). All the cell types, including Distal-Epi cells (AE progenitor cells), were detected at similar frequencies, and no notable differences were observed between HT^mat^ and KO^mat^ fetuses (Figure 3B). Of note, analysis of differentially expressed genes (DEGs) in the Distal-Epi population revealed that genes involved in differentiated AE cells (e.g., *Sftpc*, *Sftpb*, *Aqp5*) were rather highly upregulated in KO^mat^ fetuses (Figure 3C and Supplementary Table 2), even though their protein levels remained lower in the lungs of *Cyp27a1* KO^mat^ fetuses (Figure 1E), suggesting the occurrence of a translational failure in the cells. Several ribosome genes, such as *Rps29*, *Rps26*, *Rps15*, *Rpl36*, and *Rpl41*, were also highly upregulated in the Distal-Epi of *Cyp27a1* KO^mat^ fetuses (Figure 3C and Supplementary Table 2). However, similar to Sftpc and Aqp5, the corresponding ribosomal proteins were expressed at similar or lower levels (Figure 3D and Supplementary Figure S5A).

**Figure 3.**
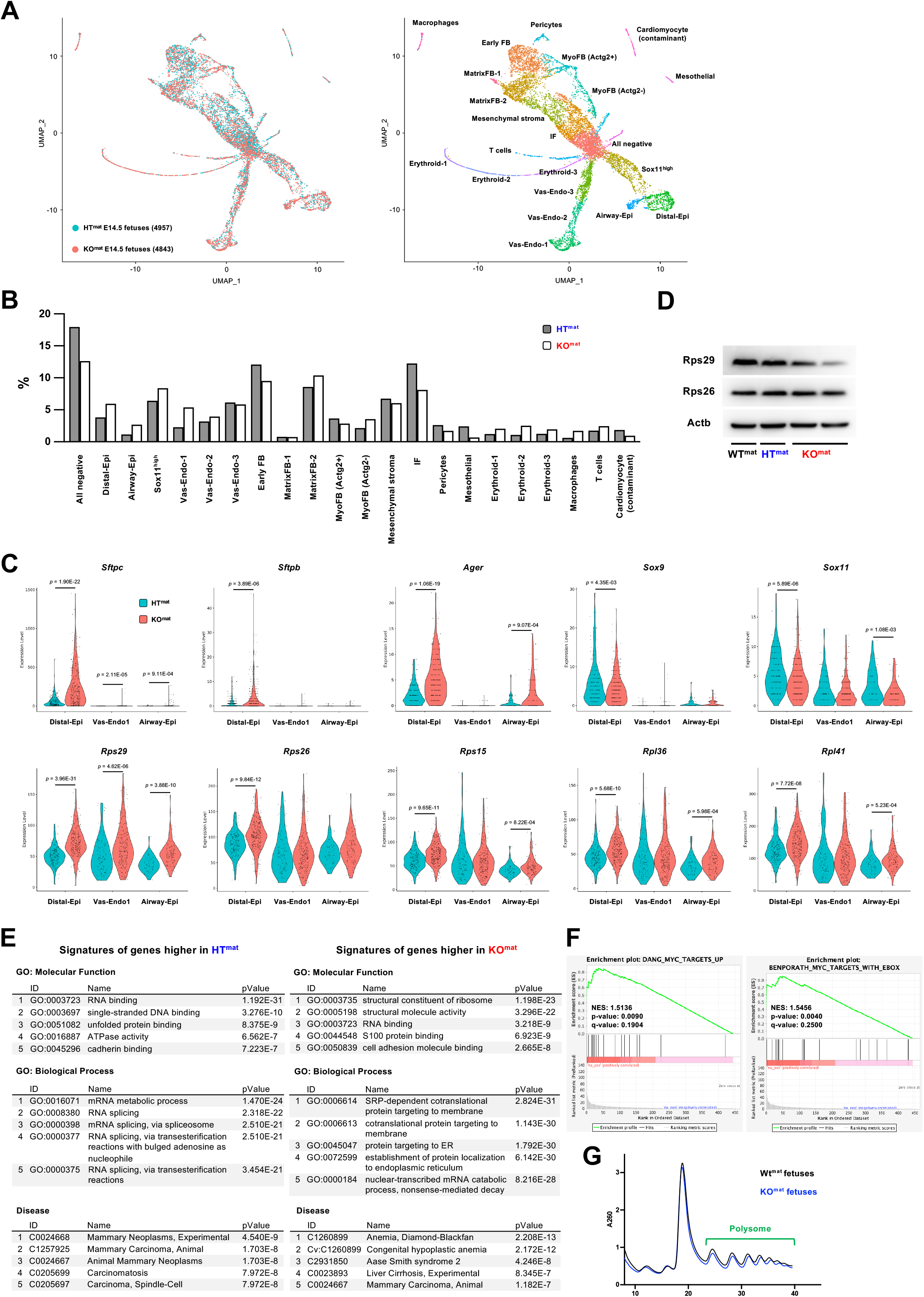
Low polysome formation in lung cells of *Cyp27a1* KO^mat^ fetuses. **(A)** scRNA-seq analysis of lung tissues from *Cyp27a^-^* HT^mat^ and KO^mat^ KO embryos. UMAP plots of the pool of cells from three embryos are shown. **(B)** Frequencies of each cluster of *Cyp27a^-^* HT^mat^ and KO^mat^ KO embryos. **(C)** Violin plots of representative genes significantly differentially expressed in Distal-Epi, Vas-Endo-1. Or Airway-Epi clusters. **(D)** Western blot analyses of Rps29, Rps26, and Actb in the lung tissues of fetuses delivered from different mothers. **(E)** GO terms related to the genes differentially expressed in Distal-Epi cells. **(F)** GSEA plots of genes significantly up-regulated in Distal-Epi cells of KO^mat^ KO embryos. **(G)** Polysome profiling of the lungs of WT^mat^ and KO^mat^ fetuses. \**p* < 0.05, \*\**p* < 0.01, \*\*\**p* < 0.001.

Gene ontology (GO) analysis using ToppFun also revealed that the genes upregulated in KO^mat^ fetal lung cells were mainly implicated in ribosome biogenesis and the translocation of proteins to the ER (Figure 3E). BAs function as a chemical chaperone that inhibits the induction of ER stress, both *in vitro* and *in vivo* (Özcan et al., 2011; Miharada et al., 2014; Sigurdsson et al., 2016; Sigurdsson et al., 2020; Koide et al., 2022). However, we failed to observe any induction of ER stress markers (Supplementary Figure S5B), suggesting the involvement of a different mechanism. In contrast, the genes that were significantly upregulated in the HT^mat^ (downregulated in the KO^mat^) fetuses were mainly involved in RNA processing (Figure 3E). Furthermore, gene set enrichment analysis (GSEA; Subramanian et al., 2005) showed that genes expressed at higher levels in the Distal-Epi of HT^mat^ fetuses than in KO^mat^ fetuses were enriched for the Myc signal, which governs translation control by regulating ribosome biogenesis (van Riggelen et al., 2010) (Figure 3F). These results imply that protein synthesis is affected in KO^mat^ fetuses. The Disease category of GO analysis indicated that genes altered in KO^mat^ fetal lung cells are implicated in Diamond-Blackfan Anemia (DBA) (Figure 3E). DBA is a congenital disease caused by a mutation or deletion in one of the ribosomal genes or *Gata1* gene, resulting in anemia due to abnormal erythroid differentiation (Da Costa et al., 2020). Defects in *Rps29* and *Rps26* are also known to cause DBA (Doherty et al., 2010; Mirabello et al., 2014). Interestingly, we found that erythroid cells in the liver of *Cyp27a1* KO^mat^ fetuses showed an abnormal differentiation profile, similar to that in DBA mouse models (Jaako et al., 2011) (Supplementary Figures S5C-E). Since patients with DBA and animal models usually show lower levels of polysome formation (Horos et al., 2012), and failures in ribosome assembly are known to cause feedback signals (Gómez-Herreros et al., 2017; de la Cruz et al., 2018), we analyzed polysome formations in the lungs of *Cyp27a1* KO^mat^ fetuses using polysome profiling (Gandin et al., 2014; Liang et al., 2018). We found that fewer polysomes were formed in the lungs of *Cyp27a1* KO^mat^ fetuses (Figure 3F), confirming the scRNA-seq observations.

### 7α-HC causes lung abnormality in *Cyp27a1* KO^mat^ fetuses

To uncover the molecular mechanisms underlying the abnormalities in *Cyp27a1* KO^mat^ fetuses, we administered BAs to these mice. Administration of 27-HC and BAs, especially 3β-Hydroxycholest-5-en-(25R)26-oic acid (Cholestenoic acid), 7α-Hydroxy-3-oxocholest-4-en-(25R)26-oic acid (7-HOCA), and taurocholic acid (TCA), which show decreased levels in *Cyp27a1^-/-^*mice (Sigurdsson et al., 2016; Griffiths et al., 2019) (Supplementary Figure S6A), did not rescue the RDS phenotype (data not shown). We observed that the concentrations of cholestenoic acid and 7-HOCA were not decreased in the livers of *Cyp27a1* KO^mat^ fetuses but were significantly lower in the blood plasma of *Cyp27a1^-/-^* adult mice (Supplementary Figure S6A and B). These observations suggested that the lung abnormalities were not due to decreased 27-HC or BA levels. It has been reported that some of the intermediate products of BA synthesis, oxysterols, are highly accumulated in *Cyp27a1^-/-^* mice (Griffiths et al., 2019). We confirmed increased oxysterols in the livers of *Cyp27a1* KO^mat^ fetuses, similar to the plasma of adult *Cyp27a1^-/-^* mice (Figure 4A and Supplementary Figure S6C). Therefore, we hypothesized that the increased oxysterol levels might cause abnormalities in the KO^mat^ fetuses. To this end, we intraperitoneally injected two major oxysterols, 7α-HC and C4, accumulating in *Cyp27a1^-/-^* mice (Figure 4A and Supplementary Figure S6C) into WT pregnant mice (Figure 4B) and found that administering 7α-HC caused a certain number of fetuses to die during pregnancy or develop RDS after birth (Figure 4C and Supplementary Movie 4). These newborns had stomachs filled with air, and the lungs could not float, as observed in *Cyp27a1* KO^mat^ fetuses (Figure 4D). The low frequency of dead/RDS offspring upon 7α-HC administration was presumably because the WT mice efficiently degraded 7α-HC through their normal Cyp27a1 activity. To clarify whether 7α-HC directly affects fetal lung maturation, we extracted WT fetal lungs and performed *in vitro* whole tissue cultures using an air-liquid interface system (Carraro et al., 2010), which enables natural alveolar branching (Figure 4E). We observed that adding 7α-HC reduced the branching of the bronchial tubes (Figures 4F and G and Supplementary Figure S7A). Furthermore, western blot analyses showed that the up-regulation of Sftpc expression upon the *in vitro* culture was suppressed by the 7α-HC addition (Figures 4H and I and Supplementary Figure S7B). We also analyzed mice lacking another key enzyme in the alternative pathway, oxysterol 7α-hydroxylase (Cyp7b1), which exhibits elevated levels of various oxysterols but not 7α-HC (Li-Hawkins et al., 2000; Meljon et al., 2019). We established different mating combinations; however, *Cyp7b1* KO^mat^ newborns were delivered without noticeable abnormalities and with the same litter size (Supplementary Figure S7C). These results indicate that 7α-HC is one of the critical metabolites creating the abnormalities seen in the *Cyp27a1* KO^mat^ fetuses, and at least partially, directly inhibiting lung development.

**Figure 4.**
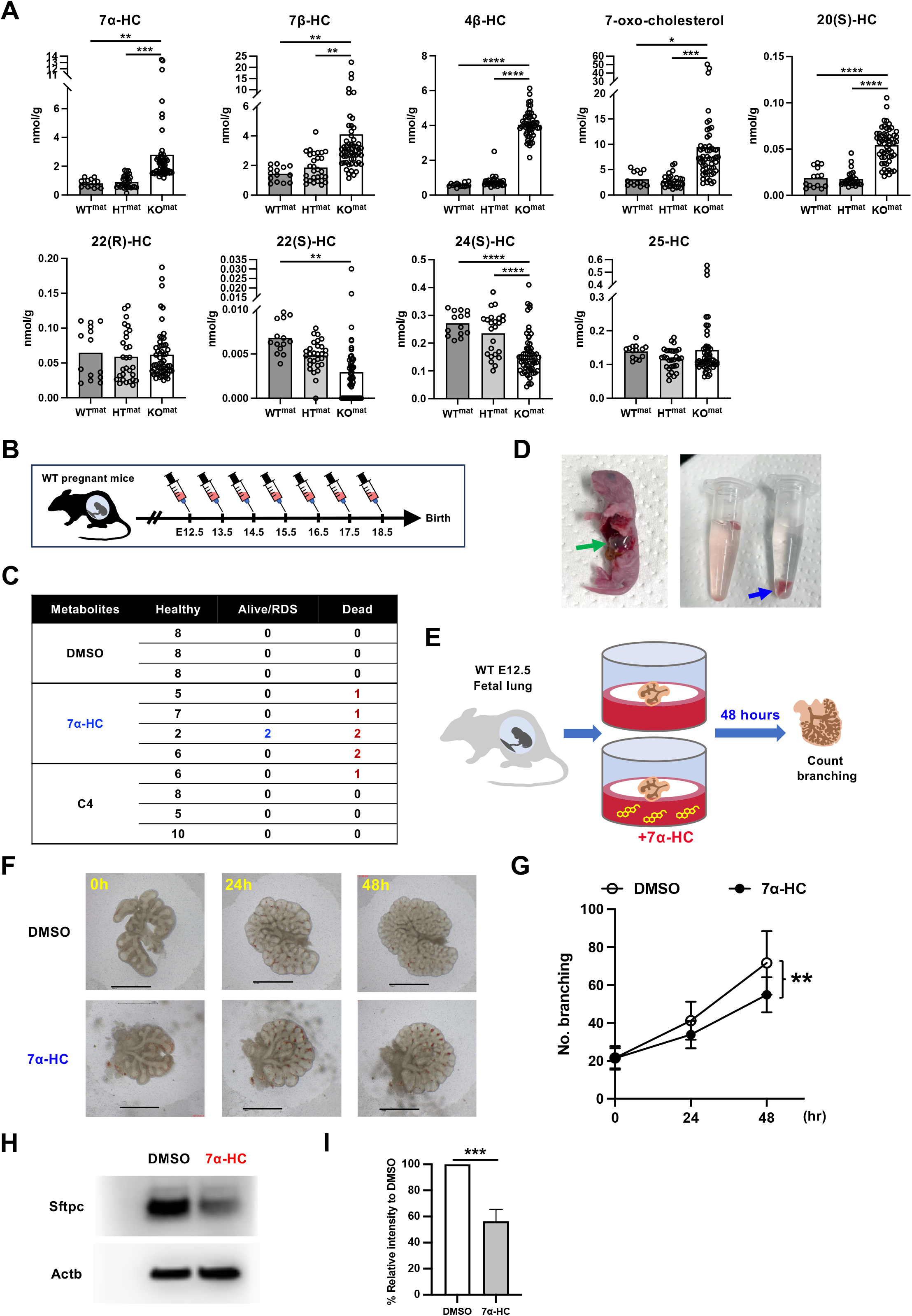
7α-HC causes the lung abnormality in *Cyp27a1* KO^mat^ fetuses. **(A)** Alterations of the oxysterol concentration in the fetal lungs. Concentrations of the following oxysterols in WT^mat^, HT^mat^, and KO^mat^ are shown; 7α-hydroxycholesterol (7α-HC), 7β-hydroxycholesterol (7β-HC), 4β-hydroxycholesterol (4β-HC), 7-oxo-cholesterol, 20(S)-hydroxycholesterol (20(S)-HC), 22(R)-hydroxycholesterol (22(R)-HC), 22(S)-hydroxycholesterol (22(S)-HC), 24(S)-hydroxycholesterol (24(S)-HC), 25-hydroxycholesterol (25-HC). **(B)** The time schedule of the injection experiment. DMSO control, 7α-HC, or 7α-Hydroxycholest-4-en-3-one (C4) was injected daily from E12.5 to E18.5. **(C)** The outcome of the injection experiment. **(D)** Representative photos of a newborn mouse derived from the mother injected with 7α-HC. Left: the green arrow indicates the stomach filled with air. Right: the blue arrow indicates the lung which is not buoyant in PBS. **(E)** A schema of the *in vitro* fetal lung culture using the air-liquid interface system. Fetal lungs extracted from WT E12.5 fetuses were placed onto Nuclepore™ for 48 hours with supplementation of DMSO or 7α-HC. **(F)** Representative photos of the fetal lungs cultured in the air-liquid interface system. The scale bar represents 1 mm. **(G)** The branching of the alveolar. The number of branching points was counted on the photos taken under the microscope. Please also see Supplementary Figure S6B. **(H)** Representative Western blot analysis of the fetal lungs cultured with or without 7α-HC. **(I)** The relative signal of Sftpc. The intensity of the Sftpc band was normalized using Actb, and the fold change in the 7α-HC samples compared to the DMSO control is shown. \**p* < 0.05, \*\**p* < 0.01, \*\*\**p* < 0.001, \*\*\*\**p* < 0.0001.

### 7α-HC targets the ribosomal protein Fau

Oxysterols bind to oxysterol-binding proteins, resulting in various biological activities (Hammond and Pacheco, 2019). 7α-HC is synthesized from cholesterol by Cyp7a1, the starting enzyme of the classic pathway (Johnson and Lack, 1976); however, the target of 7α-HC and its physiological function remain unclear. Therefore, we employed an advanced proteomics technology, Proteome Integral Solubility Alterations (PISA) (Gaetani et al., 2019), to identify 7α-HC target proteins. PISA relies on the fact that the conformational changes upon ligand binding can affect the thermal stability of proteins; therefore, whether the total amount of unprecipitated protein after heating is changed is used as a judgment criterion for ligand binding. Lysate was prepared from the E12.5 fetal lung of WT mice, and candidate chemical compounds (7α-HC, 7-HOCA, Cholestenoic acid) were compared (Figure 5A). We detected 7,010 proteins; among them, 193 proteins showed changes in quantity upon adding 7α-HC, suggesting a potential targeting (Figures 5B and C). The protein that showed the most significant change in quantity was Fau (Figure 5C and Supplementary Table 3). *Fau* encodes a fusion protein of a ribosomal protein, eS30, and a ubiquitin-like protein, Fubi (also known as monoclonal non-specific suppressor factor; Mnsfβ), which regulates the assembly of ribosomal proteins (van den Heuvel et al., 2021). In similar to *Rps29* and *Rps26*, *Fau* mRNA level was also significantly up-regulated in the Distal-Epi cells of Cyp27a1 KO^mat^ fetuses (Figure 5D). To confirm the effect of 7α-HC addition on the thermal stability of Fau, we separately analyzed the thermal stability of Fau protein upon heating, as previously employed in the CETSA (Martinez Molina et al., 2013). We utilized the murine lung carcinoma cell line, LLC (Bertram and Janik, 1980). After high-speed centrifugation to remove the insoluble fraction of the heated mixture of the LLC lysate and 7α-HC, each fraction was subjected to western blot analyses. The assay showed that at lower temperatures, the intensity of Fau mixed with 7α-HC became weaker than that without 7α-HC, thus confirming the PISA results (Figures 5E and F). Since the scRNA-seq of E14.5 fetal lungs indicated ribosomal defects in KO^mat^ fetuses and the polysomal profiling showed impaired polysome formation (Figure 3G), we tested whether 7α-HC treatment affects polysome formation. Polysome profiling revealed that the addition of 7α-HC also resulted in lower polysome formation in LLC cells (Figure 5G), suggesting that 7α-HC targets *Fau* and inhibits its function in the ribosomal assembly. These results propose that 7α-HC targets the ribosomal protein Fau and impairs polysome formation in the *Cyp27a1* KO^mat^ fetuses.

**Figure 5.**
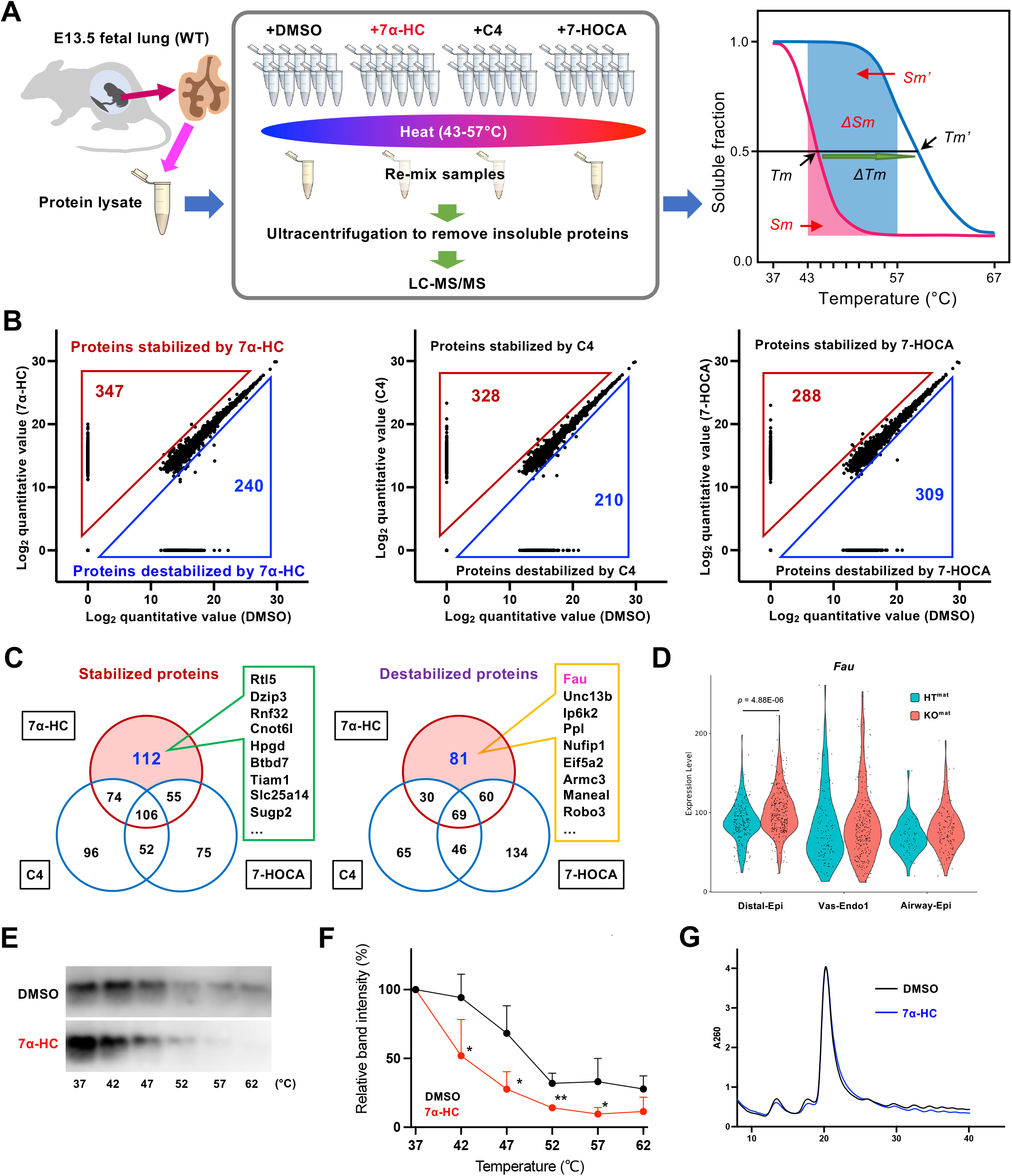
7α-HC targets a ribosomal protein Fau. **(A)** The overview of PISA on the fetal lung. Binding targets of 7α-HC and two bile acids (7-HOCA, Cholestenoic acid) were searched. The difference of the remaining soluble fraction (Δ*Sm*) indicates a thermal stability change. **(B)** Results of PISA. Log_2_ quantitative values of each protein are shown. **(C)** Venn diagrams showing overlapping stabilized/destabilized proteins between DMSO and candidate compounds. Above 1.2-fold or below 0.83-fold changes were considered meaningful differences. The top 10 proteins uniquely stabilized/destabilized by 7α-HC are listed aside from the diagrams. Please also see Supplementary Table 2. **(D)** The mRNA expression levels of *Fau* in Distal-Epi, Vas-Endo-1. Or Airway-Epi clusters. **(E)** Validation of the effect of 7α-HC on the thermal stability of Fau protein. A representative western blot analysis of the soluble fractions of LLC lysate after heating at each temperature is shown. **(F)** A summary of the western blot analyses. Band intensities relative to the 37°C sample are shown. The Fau band intensities were normalized by Actb band intensities. n=4. **(G)** Polysome profiling of LLC cells treated with DMSO or 7α-HC. \**p* < 0.05, \*\**p* < 0.01.

### Reduced *Fau* expression inhibits fetal lung maturation

To demonstrate whether reduced Fau function affects fetal lung maturation, we used shRNA to knock down *Fau* expression in fetal lungs using lentiviruses expressing shRNA against *Fau*. First, we confirmed the *Fau* down-regulation efficiency of the shRNA constructs (Figure 6A). We found the successful down-regulation of *Fau* (shFau-3 and -4) led to lower protein synthesis in LLC cells, as determined via OP-Puro staining (Liu et al., 2012) (Figure 6B). Furthermore, shFau-expressing LLC cells exhibited reduced polysome formation (Figure 6C). To down-regulate the *Fau* expression in fetal lungs, we dropped concentrated lentiviruses directly onto isolated E12.5 fetal lungs and cultured them in the ALI culture system (Figure 6D). The fetal lungs transduced with the shFau lentiviruses showed impaired alveolar branching similar to that observed upon the addition of 7α-HC (Figures 6E-F). Furthermore, the expression of the representative alveolar differentiation marker Sftpc was significantly reduced (Figures 6G and H and Supplementary Figure S7D), indicating that polysome formation mediated by Fau is necessary for alveolar maturation.

**Figure 6.**
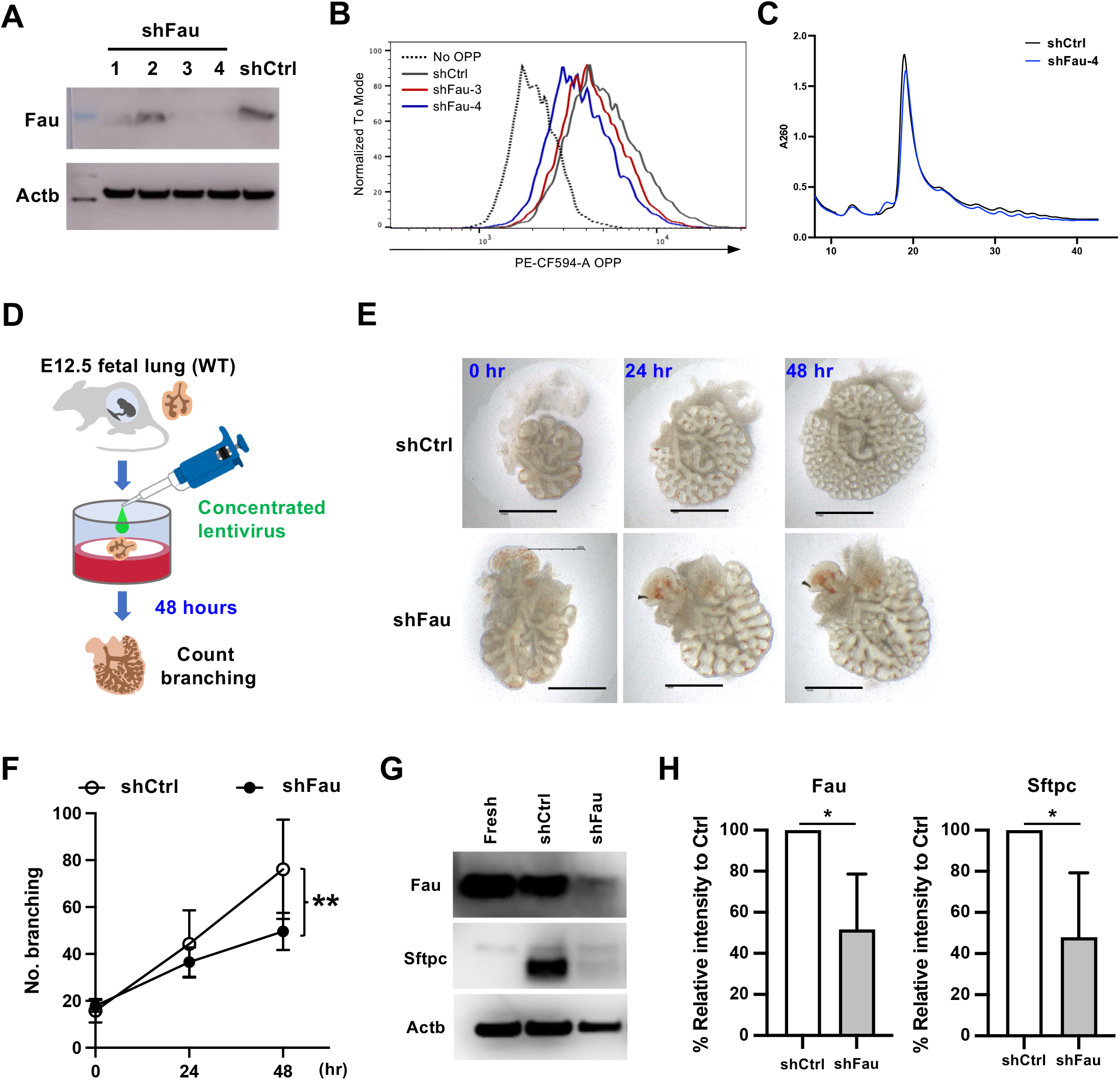
Reduced *Fau* expression inhibits fetal lung maturation. **(A)** Western blot analysis to validate the knockdown efficiency of shRNA against *Fau* (shFau). Non-targeting control shRNA (shCtrl) was used as a control. **(B)** Protein synthesis rate analysis using OP-Puro staining. A representative flow cytometry analysis of OP-Puro intensity in the LLC expressing shCtrl or shFau is shown. **(C)** Polysome profiling of LLC expressing shFau driven by the Tet-On inducible system. **(D)** A schema of the *in vitro* culture of fetal lung expressing shFau. **(E)** Representative photos of the fetal lungs cultured in the air-liquid interface system. The scale bar represents 1 mm. **(F)** The number of branching points in the fetal lungs transduced with lentiviruses. **(G)** Representative Western blot analysis of the fetal lungs transduced with lentiviruses. **(H)** The relative signal of Fau and Sftpc. The intensity of the band was normalized using Actb, and the fold changes in the shFau samples compared to the DMSO control is shown. \**p* < 0.05, \*\**p* < 0.01.

### 7α-HC impairs the maturation of placenta-consisting cells

Fau/Mnsfβ is highly and broadly expressed in both murine and human placenta, and *Fau*-deficient mice exhibit placental malformation (Gu et al., 2015; Yang et al., 2021). Moreover, a large cohort study has identified a single nucleotide polymorphism (SNP) in *FAU* to correlate with recurrent pregnancy loss in a Chinese Han population (Gu et al., 2018). Since 7α-HC targets Fau protein, we analyzed the placentas of *Cyp27a1* KO^mat^ fetuses. The placentas from *Cyp27a1* HT^mat^ and *Cyp27a1* KO^mat^ fetuses appeared normal (Figure 7A); however, their diameters and weights were significantly smaller than those from WT mothers (Figures 7B and C). Furthermore, histological analyses using H&E staining and immunostaining for cytokeratin 7 (CK7) (Yang et al., 2021) revealed that the labyrinth area of the placentas of *Cyp27a1* KO^mat^ fetuses was significantly thinner than that of *Cyp27a1* HT^mat^ and WT fetuses (Figures 7D-F), as reported in the *Fau*-deficient mice (Gu et al., 2015; Yang et al., 2021). Similar to that in the fetal lung, polysome formation in the placenta of *Cyp27a1* KO^mat^ fetuses was impaired (Figure 7G), suggesting an abnormality in translation. To clarify whether the maturation of the labyrinth area was also affected, the expression levels of syncytiotrophoblast (SynT) markers (Mct1 and Mct4) and sinusoidal trophoblast giant cell (sTGC) markers (Ctsq) (Nagai et al., 2010; Chhabra et al., 2012) were analyzed using western blot. We found that the expression of a representative marker of type I SynT (SynT-I), Mct1, was lower in the placentas of *Cyp27a1* KO^mat^ fetuses than that of WT^mat^ fetuses (Figures 7H and I and Supplementary Figure S7E). The expression of Mct1 is restricted to the apical plasma membrane area of SynT-I, which faces the maternal circulation (Nagai et al., 2010; Ueno et al., 2013). Immunostaining revealed Mct1 as thick lines expression in the placentas of *Cyp27a1* HT^mat^ and WT^mat^ fetuses. In contrast, *Cyp27a1* KO^mat^ placentas showed that Mct1 expression was weaker in the membrane area and mislocated in the cytosol (Figure 7J), indicating abnormal cell differentiation.

**Figure 7.**
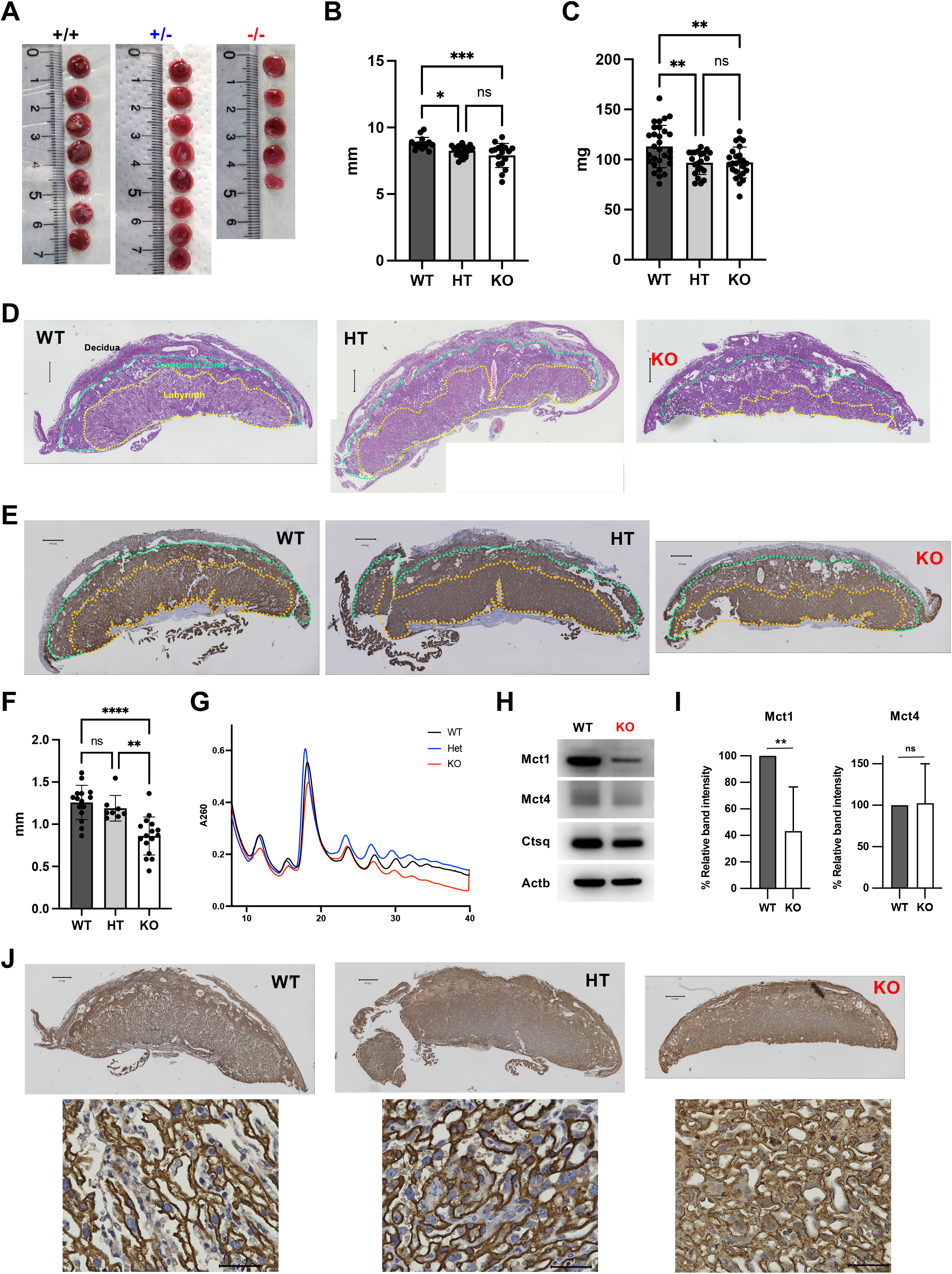
7α-HC impairs the maturation of placenta-consisting cells. **(A)** Photos of E16.5 placentas from each genotype. **(B)** The diameter of the placenta. **(C)** The weight of the placenta. **(D)** H&E staining of the placentas. The scale bar represents 500 μm. **(E)** Immunohistochemistry staining for cytokeratin 7 (CK7) of the placentas. The scale bar represents 500 μm. **(F)** Thickness of the labyrinth area. The thickness was measured on the H&E staining samples. **(G)** Polysome profiling of the placentas. **(H)** Representative western blot analyses for Mct1 and Mct4 (SynT marker) and Ctsq (sTGC marker). **(I)** The relative signal of Mct1 and Mct4. The intensity of the band was normalized using Actb, and the fold changes in the KO samples compared to the WT control are shown. **(J)** Immunohistochemistry of Mct1 in the placenta. The scale bar represents 500 μm (smaller magnification) and 50 μm (larger magnification). \**p* < 0.05, \*\**p* < 0.01, \*\*\**p* < 0.001, \*\*\*\**p* < 0.0001.

## DISCUSSION

Recurrent pregnancy loss (also known as repeated miscarriages or recurrent spontaneous abortion) is a common medical condition in many countries, as 1 - 5 % of women trying to conceive have experienced it (Baek et al., 2007; Hong Li and Marren, 2018; Dimitriadis et al., 2020; Deng et al., 2022). Even though the key parameters, e.g., embryonic aneuploidy, uterine abnormalities, autoimmune disorders, and vitamin D deficiency (Dimitriadis et al., 2020), have been discovered to cause recurrent pregnancy loss potentially, the etiology remains unknown in approximately 30-50 % of cases depending on the age (Tur-Torres et al., 2017; Deng et al., 2022). Detoxification by the maternal body is as important as providing the nutrients and metabolites required during pregnancy. In this study, we discovered an essential role of maternal Cyp27a1 in protecting developing fetuses from toxicity oriented from the excessive 7α-HC. Pregnant *Cyp27a1^-/-^* mice presented multiple abnormalities, including a low fertility ratio, smaller litter size, high stillbirth rate, and lung malformations in the delivered newborns (Figure 1B). Interestingly, all these phenotypes depended entirely on the mother’s genotype but were not influenced by fetal genotypes. Considering their timing, these events may have occurred independently of each other. The uteri of E14.5 *Cyp27a1^-/-^* mothers often showed traces of absorbed embryos, indicating that fetal growth was terminated in the early phase. We observed two types of fetuses in *Cyp27a1^-/-^* mothers later than E16.5, with and without apparent abnormalities in their appearance (Figure 2A). It is noteworthy that even if the fetuses appeared normal, they were stillbirths or developed RDS (Figure 1B). Presumably, fetuses that were less affected by maternal Cyp27a1 deficiency could survive the first two waves of life-threatening events, yet malformation of the lung alveoli was unavoidable. We did not find a reason for the difference in the actual phenotypes; sex and fetal genotypes were not correlated with the outcomes (Figure 1B and Supplementary Figure S1B).

Disruption of maternal *Cyp27a1* led to the accumulation of various oxysterols, including 7α-HC, in mothers and fetuses (Figure 4A and Supplementary Figure S6C; Griffiths et al., 2019), presumably due to the metabolic capacity of the classic pathway being exceeded by the influx of more cholesterol, as the activity of the classic pathway is low in fetuses and newborns (Motta et al., 2003; Mercer et al., 2018; Memon et al., 2022). Notably, BA supplementation of the most common BAs in mice, taurocholic acid (TCA), to the *Cyp27a1^-/-^* mothers did not rescue the phenotype of *Cyp27a1* KO^mat^ newborns (data not shown). These observations suggest that one of the major roles of the enhanced alternative pathway activity might be to prevent the accumulation of specific oxysterols, i.e., 7α-HC, rather than providing additional BAs, as reported previously. A similar concept was also proposed by recent studies (Kakiyama et al., 2020; Wang et al., 2021). The deletion of Cyp27a1, which resides in mitochondria (Lorbek et al., 2011), led to an enhanced autooxidation, increasing 7-oxo-cholesterol (7-ketocholesterol) and 7β-HC (Figure 4A and Supplementary Figure S6C), which are potent apoptosis triggers (Berthier et al., 2005; Nury et al., 2021). However, lung epithelial cells of *Cyp27a1* KO^mat^ fetuses did not exhibit an increase in the number of apoptotic cells (Figures 2B and C). Therefore, 7α-HC served as an inhibitory factor, with a mechanism distinct from the previously known cytotoxicity of cholesterol/oxysterol/BA (Figures 5E-G). Thus, the inhibitory effect of 7α-HC on the ribosomal assembly might have potential adverse regulatory effects in fetal development.

In *Cyp27a1* KO^mat^ fetuses, the number of alveolar epithelial cells significantly decreased, whereas progenitors and airway epithelial cells were unaffected (Figures 1F and G). Notably, the expression of genes crucial for alveolar differentiation, *Sftpc* and *Aqp5*, was significantly higher in KO^mat^ fetuses at the transcriptional level but not at the protein level (Figures 1E and 3C). We found that 7α-HC modifies ribosome assembly by targeting the Fau protein, leading to impaired polysome formation, thereby decreasing the protein synthesis (Figures 5C, 5G, 6B, and 6C). It is not completely clear why specific cell types, such as alveolar epithelial cells, in the lungs are affected; however, one potential mechanism might involve different translation requirements. While DBA results from a deficiency or mutation of ribosomal genes ubiquitously expressed in all cells, the abnormality is limited to specific tissues or cell types, especially erythroid cells (Sankaran and Weiss, 2015). These cells require a high translation rate for the master regulator of erythroid commitment, Gata1, the translation of which is tightly and selectively regulated at the ribosome level (Khajuria et al., 2018). This machinery has been considered general over tissue/cell type (Buszczak et al., 2014). In our study, multiple essential genes for alveolar differentiation and maturation, such as *Sftpc* and *Aqp5*, showed drastic changes in the expression; their protein levels were severely reduced, while their transcription levels were highly up-regulated (Figures 1D, 1E, and 3C). In addition, DBA cells exhibit cell cycle arrest through p53 induction (Jaako et al., 2011; Le Goff et al., 2021). We also found that EpCAM^+^ epithelial cells in *Cyp27a1* KO^mat^ fetuses stopped proliferating (Figures 2D and E), although no change in p53 levels were observed (data not shown). These multilayered aberrations are considered to be the cause of the alveolar malformations. Of note, concentrations of specific oxysterols, including 7α-HC, are elevated in *Cyp27a1* KO mice even without pregnancy (Supplementary Figure S6C and Griffiths et al., 2019); however, both male and female mice did not develop any severe phenotype unless a high-cholesterol diet was given (Rosen 1998; Sigurdsson et al., 2016). One possible reason for this is the Fau expression pattern, which is high in the placenta (Gu et al., 2015; Yang et al., 2021); therefore, its impact is limited to the fetal stage.

The ribosomal defect in the *Cyp27a1* KO^mat^ fetuses was not a result of genetic abnormalities in ribosomal genes but was caused by a metabolite interaction. Our study showed the potentially harmful effects of maternal 7α-HC. Since oxysterol production could be affected by cholesterol intake (Brooks et al., 2017; Sozen et al., 2018), developmental defects could occur without genetic mutations; i.e., overnutrition during pregnancy, such as metabolic associated steatotic liver disease (MAFLD) (Nakanishi et al., 2022; Lee et al., 2023). Our findings suggest that organ malformations and developmental failures do not necessarily involve fetal congenital abnormalities but could be triggered by environmental factors from the maternal circulations. The placentas functions as effective barriers limiting the transmission of most proteins; therefore, maternal metabolites likely play a central role as toxic factors affecting embryonic development.

*CYP27A1* deficiency in humans causes an autosomal recessive lipid storage disorder called cerebrotendinous xanthomatosis (CTX) (Björkhem, 2013). Since CTX is a rare disease, whether the deletion of *CYP27A1* could lead to pregnancy pathologies and is the primary etiology of recurrent pregnancy loss remains unclear. A study has reported a rapid increase in 27-HC levels during the first trimester of pregnancy; however, no significant differences were observed in the patients with pregnancy pathologies, including fetal growth retardation, preeclampsia, HELLP syndrome, and intrahepatic cholestasis of pregnancy (ICP) (Winkler et al., 2017). ICP often results in severe adverse perinatal outcomes, including preterm birth, intrauterine fetal demise, and stillbirth, as well as lung immaturity, and neonatal RDS (Zecca et al., 2008; Williamson and Geenes, 2014). Long-term exposure of fetuses to elevated levels of BAs can cause ICP-oriented lung immaturity and injury (Zhang et al., 2015; Futterman et al., 2022). Interestingly, our study showed that not the lack of 27-HC but rather the increased 7α-HC levels might be responsible for the fetal abnormalities. These observations imply that the adverse outcomes in fetuses of mothers with impaired cholesterol metabolism, like those with ICP, might result from the elevated levels of 7α-HC, not BAs.

In conclusion, our findings suggest that maintaining the integrity of maternal BA synthesis, particularly 27-hydroxylase activity, is essential for fetal organ formation and maturation. We propose that specific metabolites produced under altered metabolic conditions in the maternal body could critically affect embryonic development through interactions with key proteins that regulate fate commitment, differentiation, and maturation of the cells in embryos/fetuses. Further studies on altered maternal metabolic conditions would help better understand several developmental defects.

## Materials and methods

### Mice

B6.129-*Cyp27a1^tm1Elt^*/J (*Cyp27a1* KO) mice and B6;129S-*Cyp7b1^tm1Rus^*/J (*Cyp7b1* KO) were obtained from The Jackson Laboratory. Fetuses were collected from timed pregnant mice obtained from in-house breeding at the animal facility of Lund University or Kumamoto University. All animal experiments were approved by either the Lund University Animal Ethical Committee or the Kumamoto University Animal Ethical Committee.

### *In vitro* fertilization

*In vitro* fertilization of Cyp27a1 KO mice was performed by following Takeo et al., 2008 and 2010 at Lund Transgenic Core Facility.

### Sample preparation for oxysterol measurements

Extraction of oxysterols from fetal liver was performed according to the previous method (Kakiyama et al., 2020) with modifications: E14.5-18.5 mouse fetal liver (20 mg) was homogenized with *d6*-25-HC (100 pmol/ml methanol, 10 µl), butylated hydroxytoluene (5 mg/ml ethanol, 10μl) and acetonitrile/methanol, 50/50, v/v (5 ml) using NISSEI bio-mixer (Nihonseikaki Kaisha Ltd., Tokyo Japan) at 6,000 rpm for 5 minutes. After centrifugation at 1,000 *x g* for 10 minutes, the supernatant was collected in a glass tube which was then concentrated to *ca.* 0.5 ml under a nitrogen stream. Water (1 ml) and *n*-hexane (2 ml) were added, and the mixture was thoroughly vortexed for 2 minutes. Subsequently, the layer was separated by centrifugation at 1,000 *x g* for 5 min. The upper layer was collected in a glass tube. The bottom layer was washed again with *n*-hexane (1.0 ml) by the same procedure, and the combined extract was dried up under a reduced pressure.

Dried blood spot (DBS) was used for the oxysterol measurement of adult mouse plasms. A 6.0-mm-diameter disc from the center of the spot (equivalent to 5.1 µl blood with 55% hematocrit value) was placed in a 2 ml glass tube, and *d6*-25-HC (100 pmol/ml in methanol, 10 µl), butylated hydroxytoluene (5 mg/ml in ethanol, 10 µl), and water:methanol, 50:15, v/v, (0.5 ml) were successively dropped onto the spot. After letting it at room temperature for 30 min, the wet disc was ultrasonicated for 15 minutes. To extract oxysterols, methyl tert-butyl ester (MTBE) (2 ml) was added and vortexed thoroughly for 30 seconds. The layer was then separated by centrifugation at 1,000 *x g* for 10 minutes, and the supernatant was collected in a glass tube. This procedure was repeated twice, and the combined extract was dried up under a nitrogen stream.

The extract from fetal liver tissue or DBS was redissolved in *n-*hexane (0.5 ml) and loaded onto an InertSep NH2 cartridge (100 mg/1 ml; GL Sciences Inc., Tokyo, Japan) to remove excessive cholesterol. Of note, prior to loading the sample, the column was rinsed with chloroform/methanol, 1:1, v/v (1 ml), and conditioned with *n-*hexane (3 ml). After loading the sample, the column was washed with *n-*hexane (1 ml), and the desired oxysterols were eluted with chloroform/methanol, 20:1, v/v (1 ml). Oxysterols were then derivatized to nicotinyl ester form as follows (Sidhu et al., 2015): the derivatization reagent (100 µl), which was a mixture of nicotinic acid (80 mg), N,N′-dimethyl-4-aminopyridine (30 mg), and 1-ethyl-3-(3-dimethylaminopropyl) carbodiimide hydrochloride (100 mg) in N,N′-dimethylformamide (1.0 ml), was added to the oxysterol sample, and the mixture was incubated at 60°C for 1 h. Water (0.5 ml) and *n-*hexane (1.0 ml) were added, and the solution was thoroughly vortexed for 2 minutes. After centrifugation at 180 *x g* for 5 min, the *n-*hexane layer was evaporated under a nitrogen stream. The residue was re-dissolved in acetonitrile (100 µl), and an aliquot (10 µl) was injected to the liquid chromatography-mass spectrometry (LC-MS) system.

### Oxysterol measurement by LC-MS

LCMS-8050 tandem mass spectrometer equipped with an electrospray ionization (ESI) probe and Nexera X2 ultra-high-pressure LC system (Shimadzu Co., Kyoto, Japan) was used. A SunShell C30 column (100 mm × 2.1 mm inner diameter, 2.6 µm particle size; ChromaNik Tecnology Inc., Osaka, Japan) was used at 35°C for chromatographic separation. Mobile phase A (5 mM ammonium acetate containing 0.1% formic acid) and mobile phase B [isopropanol-acetonitrile (3:2, v/v) containing 0.1% formic acid] were used for the gradient elution as follows: 0–2 min 70% B; 2–6 min 70–76% B; 6–16 min 78% B; 16–18 min 96% B; 18–20 min 96% B; 20–20.1 min 96–70% B; and 20.1–25 min 70% B. Flow rate was kept constant at 0.25 ml/min. The nebulizer gas flow was set at 3.0 litter per minute, and the heating gas and the drying gas were set at 10 liters per minute. The interface temperature was 300°C; the de-solvation line temperature was 250°C; and the heat block temperature was set at 400°C with an interface voltage of 4,000 V. The collision gas (argon) pressure was 270 kPa. Quantification of all oxysterol analytes was achieved in positive multiple reaction monitoring (MRM) mode (Supplementary Table 4).

### Sample preparation for cholestenoic acids and bile acids and measurements

The concentrated supernatant from the fetal liver homogenate (See sample preparation for oxysterol measurements) was combined with 100 µl of internal standard (IS) solution (a mixture of 1.0 nmol/ml of *d4*-CA, *d4*-GCA, *d4*-TCA, *d4*-GCDCA, *d4*-TCDCA, *d5*-CDCA-3S, *d5*-GCDCA-3S, and *d5*-TCDCA-3S in 50% ethanol). The homogenate containing IS was then diluted with water (5 ml) and loaded onto a preconditioned InertSep C18-B cartridge (500 mg/6 ml). After washing the column with water (6 ml), cholestenoic acids and bile acids were eluted with 90% ethanol (4 ml). The collected effluent was dried up under a nitrogen stream, and re-dissolved in 20% acetonitrile (1 ml). A 5 µl aliquot was injected into the Shimadzu LCMS-8050 system described below.

Extraction of cholestenoic acids and bile acids from DBS was carried out essentially according to the method by Muto et al (Muto et al., 2023): A 6.0-mm-diameter-disk (equivalent to 5.1 µl blood with 55% hematocrit value) was placed in a 2-ml glass tube, and the fixing solution, methanol:acetone:water, 7:7:2, v/v/v (20 µl) was dropped onto it. After letting at room temperature for 30 min, the disc was dried up at 37°C for 30 minutes. The IS solution (a mixture of 1.0 nmol/ml *d4*-TCA, *d5*-CA, and *d5*-TCDCA-3S in methanol, 10 µl) and 90% methanol (100 µl) were dropped onto the dried disc and ultrasonicated for 15 min. After letting for 1-minute, a clear supernatant was taken to a sample vial, and 10 µl aliquot was injected into the LC-MS system described below.

### Cholestenoic acids and bile acid measurements by LC-MS

Shimadzu LCMS-8050 tandem mass spectrometer equipped with an ESI probe and Nexera X2 ultra-high-pressure LC system was used. An InertSustain C18 column (150 × 2.1 mm inner diameter, 3 µm particle size; GL Sciences) was used at 40°C for chromatographic separation. Mobile phase A (5 mM ammonium acetate) and mobile phase B (acetonitrile) were used for gradient elution as follows: 0–0.5 min 14% B; 0.5–5 min 14–22% B; 5–36 min 22–60% B; 36–46 min 60–98% B; 46–50 min 98% B; 50–50.1 min 98–14% B; and 50.1–56.0 min 14% B. Flow rate was kept constant at 0.2 ml/min. The nebulizer gas flow was set at 3.0 liters per minute. The heating gas and the drying gas were set at 10 liters per minute. The interface temperature, de-solvation line temperature, and heat block temperature were set at 300°C, 250°C, and 400°C, respectively. The interface voltage was -3,000 V. Quantitative analysis was performed using MRM transition pairs for each analyte in negative ion mode (Supplementary Table 4).

### Cell line and cell culture

LLC (RCB0558) and OP9 (RCB1124) were obtained from the RIKEN Cell Bank (Tsukuba, Ibaraki, Japan) and maintained as indicated by the distributor.

### Histology analysis (H&E staining, Immunohistochemistry)

Fetal lungs and placentas were isolated and fixed in 4% paraformaldehyde at 4°C for 16 hours, dehydrated in graded ethanol solutions, cleared in xylene, and embedded in paraffin. Paraffin-embedded samples were cut into 4 μm sections and used for histological analysis.

Hematoxylin and eosin (H&E) staining was performed using standard protocols. Briefly, the sections were cleared twice with xylene, rehydrated and equilibrated in water, and washed in PBS. The rehydrated sections were treated with a hematoxylin solution. The sections were then rinsed thoroughly with water and then treated with eosin solution. The stained sections were dehydrated with graded alcohols, washed with xylene, and mounted with coverslips using a mounting medium.

For immunohistochemistry (IHC), antigen retrieval was carried out in an autoclave using Tris-EDTA buffer (pH 9.0, for Cytokeratin 7) or 0.01 M citrate buffer (pH6.0, for Mct1). After treating with 35% H_2_O_2_ and methanol for 30 min. to block endogenous peroxidase activity, sections were incubated in Tris-buffered saline/0.1% Tween-20 (TBS-T) containing 2.5% goat serum (included in ImmPRESS HRP Goat Anti-Rabbit IgG Polymer Kit, Vector Laboratories, MP-MP-7451-15) to block non-specific binding. The following primary antibodies were used: anti-Cytokeratin 7 (1/8,000, ab181598, abcam), and - Monocarboxylic acid transporter 1 (1/1,800 (0.5 μg/ml), ab85021, abcam). ImmPRESS HRP Goat Anti-Rabbit IgG Polymer Kit was used as the secondary antibody. For visualization, ImmPACT DAB Peroxidase Substrate Kit (Vector Laboratories, SK-4105) was used. Nuclei were counterstained with hematoxylin. All slides were imaged using BZ-X800 All-in-One microscope (KEYENCE).

### Immunostaining

Total lung from the trachea was taken and fixative (4% paraformaldehyde in PBS), dehydrated in graded ethanol solutions, cleared in xylene, and embedded in paraffin. For immunostaining staining, 4-μm-thick sections were treated for 30 min at 60℃ to ensure adherence to the slides. After de-paraffinize slides as usual, a PT Link (Dako) warm bath treatment device with a stable heating function was used as the antigen activation device. Heating was started by immersing the sections in an antigen activation solution (EnVision FLEX Target Retrieval Solution, Low pH, K800521-2) preheated to 65°C. After the antigen activation solution reached 97°C, heat treatment was performed for 20 minutes. After rinsing thoroughly with PBS, the sections were treated with blocking by 2% bovine serum solution for 1 hour at room temperature to prevent non-specific binding of the primary antibody. The tissue sections were then incubated with a primary antibody specific to the target protein of interest for 1 hour at room temperature, protected from light. The tissue sections were then incubated with a secondary antibody for 45 min at room temperature. Coverslips were mounted onto glass slides with an anti-fade mounting medium with DAPI (ProLong™ Gold Antifade Mountant, Invitrogen™) and visualized with a microscope.

### Single-cell RNA sequencing analysis

Lungs extracted from E14.5 or E18.5 embryos were minced and digested using collagenase I and dispase I (both 1 mg/mL, Roche) and incubated at 37°C for 5 min. for twice. The cells were transferred to a 50 ml tube through a 100 μm cell strainer. After rinsing the cell strainer with 1 ml of PBS twice, 5 ml of PBS was added to the tube through the cell strainer. The cell suspension was transferred to a new 50 ml tube through a 40 μm cell strainer. Cells were collected by centrifugation at 200 x g for 5 min. and re-suspended with 1 ml of ammonium chloride solution for red blood cell lysis. After adding 5 ml of the staining medium, the cells were then transferred to a 15 ml tube through a 50 μm filter and centrifuged for 5 min at 200 x g.

Before sequencing, isolated single cell preparations were subjected library preparation using a Chromium system (10x Genomics). The libraries were generated according to the manufacturer’s instructions. In short; the isolated single cells were encapsulated into GEMs using the 10x genomics 3’ single cell reagent kit. Barcoded libraries were then generated within these GEMs after cell lysis and subsequent cDNA synthesis via reverse transcription. The cDNA is then amplified and subjected to fragmentation, end repair, A-tailing, adapter ligation, and sample index PCR. This process generated libraries that were then sequenced. The libraries were sequenced on Nextseq500 or Novaseq 6000. The raw cell ranger count FASTQ data from each GEM well was pooled using cell ranger aggr and analyzed according to the Cell RangerTM pipeline (10x Genomics software, version 2.0.2). All analyses, including log normalization, scaling, clustering of cells, and identifying cluster marker genes, were performed using R software package Seurat (Butler et al., 2018) version 3.1. We used the following parameters to filter out low quality cells: gene and unique molecular identifier (UMI) filtering number of UMI (nUMI), minimum 1000/maximum 75000; percent.mito, minimum 0%/maximum 5%; average number of reads/cell, 25,000. We performed principal component analysis dimensionality reduction with highly variable genes as input. Unsupervised clustering was performed using the Seurat “kmeans” function.

### Erythroid lineage analysis on flow cytometry

Fetal livers were extracted from E16.5 fetuses, and pipetted to prepare single cell suspensions. The cells were treated with an ammonium chloride solution (STEMCELL Technologies) for red blood cell lysis. After washing with PBS containing 2.5 % fetal bovine serum (FBS), the cells were stained with Ter119 antibody (BioLegend), anti-CD44 antibody (IM7, BioLegend) and 7-Amino-Actinomycin-D (7AAD, Sigma-Aldrich). Cells were analyzed on FACS LSRII (BD), and collected data were analyzed using FlowJo software (BD).

### Cell cycle assay (BrdU assay) and apoptosis assay (Annexin-V assay)

Pregnant mice were intraperitoneally injected with 120μL (1.2mg) of BrdU solution. Twenty-four hours later, the mice were sacrificed, and the fetal lungs were collected and put into PBS (Ca+, Mg+) containing 1 mg/ml collagenase type I and 1 mg/ml dispase. The tissues were minced and triturated in the solution and incubated at 37 °C for 5 min in the incubator, and the process was repeated once more. The cell suspension was then transferred into a 50 ml tube through a 70 μm cell strainer, and 8 ml of PBS was added through the cell strainer. The flow-through was transferred to a new 50 ml tube through a 40 μm cell strainer, and centrifuged at 350 xg for 5 min. After removing the supernatant, the pellet was re-suspended with 1 ml of ammonium chloride solution and incubated for 5 min at room temperature to remove red blood cells. Staining Medium (SM; PBS supplemented with 2.5 % FBS) was added to the tube to terminate the reaction, and the cell suspension was then transferred into a 15 ml tube through a 50 μm filter and centrifuged for 5 min at 350 xg. After discarding the supernatant, cells were re-suspended with 1 ml SM and the cell number was counted, and adjusted to 1×10^6^ cells/ml.

For the BrdU assay, the cells were stained with antibodies for cell surface markers (CD45, Ter119, and EpCAM), and subsequently fixed and permeabilized with BD Cytoperm/Permeabilization kit. After washing, the cells were stained with fluorescent anti-BrdU and 7-AAD (BD). For the Annexin-V assay, after cell surface staining the cells were stained with FITC-conjugated Annexin-V (BD).

### OP-Puro Incorporation assays

Protein synthesis was measured by Click-iT Plus Alexa Fluor 594 Protein Synthesis Assay Kit (Invitrogen) according to the manufacture’s protocol. Briefly, 1x10^5^ cells were plated on 24 well plate and 2 days after seeding incubated with or without Click-iT OPP Reagent for 30 min at 37℃. Collected cells were washed with PBS and fixed and permeabilized using BD Cytofix/Cytoperm Fixation/Permeabilization Kit (BD Biosciences). Cells were stained with Click-iT Plus OPP reaction cocktail for 30 min at room temperature and analyzed by FACS Symphony (BD Biosciences).

### *In vitro* fetal lung culture

*The in vitro* culture method of whole fetal lung was described by Carraro et al., 2010. Briefly, lungs were extracted from E12.5 WT fetuses in Hanks Balanced Salt Solution (HBSS) under the stereoscopic dissecting microscope. The lungs were placed onto Nuclepore™ membrane (13 mm, 8.0 μm, Cytiva) on top of the D-MEM/F-12 medium supplemented with 50 units/ml of Penicillin-Streptomycin and cultured in an incubator at 37°C. DMSO or 7α-HC was added to the medium. In the case of the shRNA transduction, concentrated lentiviral vector solutions for expressing shRNA were directly poured onto the fetal lung. The branching of the alveolar was counted on the photos taken under the microscope.

### Proteome Integral Solubility Alterations (PISA)

Lung tissues were extracted from E13.5 WT C57BL/6 embryos (from 3 pregnant mice) and put into HBSS (-) solution. After adding a protease inhibitor cocktail (Roche), the tubes were put into liquid nitrogen for 5 minutes and thawed at a 37°C water bath, and the process was repeated for 4 times more. The tubes were centrifuge at 10,000 g for 10 min at 4°C to remove debris. Clear lysate was then transferred into a new tube and adjusted to 700 μl with HBSS (-). The adjusted solution was separated to 170 μl x 4 tubes, and 1.7 μl (10 mM) of chemical compounds (7α-HC, 7-HOCA, Cholestenoic acid) and incubated at room temperature for 15 min.

After the incubation, the reaction was aliquoted to 20 μl x 8 in PCR tubes, and each tube was heated at 43, 45, 47, 49, 51, 53, 55, or 57°C for 3 min followed by 6 min incubation at 12°C using thermal cyclers. All aliquots were mixed and transferred to a Beckman 1.5 ml tube and ultracentrifuged at 150,000 xg for 30 min at 4°C using MAX-XP with TLA100.3 rotor. The supernatant was transferred to a new tube (approx. 160 μl).

Each sample was solubilized with phase transfer surfactant (PTS) buffer (Masuda et al., 2008). A total of 50 ug of each sample was transferred to a 1.5 ml tube and boiled at 95°C for 5 min. Then, the samples were reduced with 10 mM TCEP, alkylated with 20 mM iodoacetamide, and protein was purified using SP3 protocol (Hughes et al., 2019). Purified protein was digested with trypsin (protein weight: 1/50) and Lys-C (protein weight: 1/50) for 16 h at 37 °C. Peptides were desalted and purified on an C18-SCX StageTips (Adachi et al., 2016). Peptides were dried up with speedvac and solubilized with 0.1% formic acid / 2% acetonitrile.

LC-MS/MS was performed by coupling an UltiMate 3000 Nano LC system (Thermo Scientific, Bremen, Germany) and an HTC-PAL autosampler (CTC Analytics, Zwingen, Switzerland) to a Q Exactive hybrid quadrupole-Orbitrap mass spectrometer (Thermo Scientific). Peptides were delivered to an analytical column (75 μm ×30 cm, packed in-house with ReproSil-Pur C18-AQ, 1.9 μm resin, Dr. Maisch, Ammerbuch, Germany) and separated at a flow rate of 280 nL/min using a 105-min gradient from 5% to 30% of solvent B (solvent A, 0.1% FA and 2% acetonitrile; solvent B, 0.1% FA and 90% acetonitrile). The Q Exactive instrument was operated in the data-independent acquisition mode (DIA). For gas-phase fractionation (GPF)-DIA acquisition of a pooled sample for library, full mass spectra were acquired with the following parameters: a resolution of 140,000, an automatic gain control (AGC) target of 3 × 10^6^ with a maximum injection time of 200 ms, and a normalized collision energy of 27. The five GPF-DIA runs collectively covered 398-802 m/z (i.e., 398-482, 478-562, 558-642, 638-722, and 718-802 m/z). MS2 spectra were collected with the following parameters: a 2-m/z isolation window at 35,000 resolution, an AGC target of 3 × 10^6^ ions, a maximum injection time of “auto”, and normalized collision energy of 27. For the individual samples for proteome profiling acquisition, full mass spectra were acquired in the range of 398–802 m/z with the following parameters: a resolution of 140,000, an AGC target of 3 × 10^6^ with a maximum injection time of 200 ms, and a normalized collision energy of 27. MS2 spectra were collected with the following parameters: a 8-m/z isolation window at 35,000 resolution, an AGC target of 3 × 10^6^ ions, a maximum injection time of “auto”, overlapping window patterns and normalized collision energy of 27. LC-MS/MS analysis was performed in Proteobiologics Co., Ltd. (Osaka, Japan).

### Data processing and visualization

Raw MS data were processed with DIA-NN software (Ver. 1.7.12). Database searching included all entries from the mouse UniProt database and contaminant database (Tyanova et al., 2016a). This database was concatenated with one composed of all protein sequences in the reversed order. The search parameters were as follows: up to two missed cleavage sites, 7-30 peptide length, carbamidomethylation of cysteine residues (+57.021 Da) as static modifications, protein names from FASTA for implicit protein grouping, robust LC (high precision) for quantification strategy, and RT-dependent for cross-run normalization. Precursor ions were adjusted to a 1% false discovery rate (FDR).

Statistical analysis was conducted using Perseus ver. 1.6.14.0 (Tyanova et al., 2016b). For the proteome data, the quantitative ion intensities were log2-transformed and normalized by the median centering of the values of top 500 precursor in each sample. Statistically significant changes were identified by two-tailed Welch’s t test using the LFQ intensities of the two groups.

### Knockdown of Fau using shRNA

Mission lentiviral shRNA plasmids targeting Fau (shFau: TRCN0000104130, TRCN0000104131, TRCN0000104132, and TRCN0000104133), non-targeting shRNA control (shCtrl: SHC016) were purchased from Sigma-Aldrich. Tetracyclin (Tet) inducible shRNA plasmids targeting Fau (TRCN0000104133) was custom-made at Cellecta. Lentiviral particles were produced by transfecting 293T cells with the shRNA constructs and lentivirus component plasmids (pCAG-HIVgp and pCMV-VSV-G-RSV-Rev) obtained from RIKEN DNA Bank (Ibaraki, Japan).

### Polysome profiling

LLC cells were transduced with Tet-inducible Fau shRNA lentiviral vectors and stable clones were established by puromycin selection. Stable clones were plated on a poly-L-lysin coated 15 cm dish, and doxycycline (DOX, 1μg/mL, SIGMA) was added to the culture medium at next day. Forty-eight hours after the DOX addition, cells were subjected to polysome isolation according to the previously described protocol (https://www.jove.com/v/51455/polysome-fractionation-analysis-mammalian-translatomes-on-genome-wide). Briefly, cycloheximide was added to the media at 100 μg/mL for 5 min at 37℃ and 5% CO2. After washing twice with ice-cold PBS containing 100 μg/mL of Cycloheximide, cells were collected by scraper and lysed in hypotonic lysis buffer: 5mM Tris-HCI pH7.5, 2.5mM MgCl2, 1.5mM KCl and 1x Protease inhibitor cocktail, 100 μg/mL Cycloheximide, 2 mM DTT, 0.5% Triton X-100, 0.5% Sodium Deoxycholate, and 100 units RNase OUT (Invitrogen). For tissue samples, modified hypotonic lysis buffer with a 10-fold higher concentration of Cycloheximide and Recombinant RNasin Ribonuclease Inhibitor (Promega) was used instead of RNase OUT. The cytosolic lysate was layered on 10%-50% sucrose gradients and centrifuged at 246,000 g for 65 min in SW55 swinging bucket rotor (Beckman). Continuous UV absorbance was monitored with Triax flowcell monitor (BioComp).

### Western blot analyses

Proteins were extracted from cells using RIPA buffer (Thermo Fisher). Bolt Bis-Tris Plus Mini Protein Gels, 4-12% (Thermo Fisher) with Bolt MES SDS Running Buffer (Thermo Fisher) were used for the separation of proteins in the samples. The electrophoresed proteins were transferred to PVDF membranes using iBlot-2 dry blotting system (Thermo Fisher). The following primary antibodies were used: anti-Fau (13581-AP, Proteintech), -Mct1 (ab85021, Abcam), -Mct4 (22787-1-AP, Proteintech), -Sftpc (ab90716, Abcam), -Rps26 (14909, Proteintech), -Rps29 (17374, Proteintech), and -Actb (3700, Cell Signaling Technology). Horseradish peroxidase (HRP) conjugated secondary antibody was used for chemiluminescent detection. Band intensities were calculated with Image Lab software (Bio-Rad).

### Statistical analysis

Statistical significance was determined using the Bon-ferroni method for comparison of multiple groups or the two-tailed Student’s t-test for comparison of two groups. All statistical analyses were performed on Prism (Graphpad) or R with R package ggpubr.

### Author contributions

KM designed the project. M.S., S.N., H.N., and K.M. planned experiments. M.S., S.N., N.M., H.T., C.M., T.S., M.A., V.S., K.S., S.K., and K.M. performed experiments. M.v.d.G., P.P., and K.M. analyzed scRNA-seq data. G.W.T., M.M., G.K., and H.N. gave specialists’ opinions. M.S., S.N., G.K., and K.M. wrote the manuscript.

## Acknowledgements

We thank Hitoshi Niwa, Mitsuru Morimoto, and Pierre Sabatier for the scientific discussions, and Yuko Nakaishi-Fukuchi for technical support. This work was supported by Japan Society for the Promotion of Science (JSPS) KAKENHI Grant-in-Aid for Scientific Research (C) [grant number JP19K08261] (M.S.), JSPS Home-Returning Researcher Development Research [grant number JP20K23380] (K.M.), JSPS Core-to-Core Program Advanced Research Networks “Integrative approach for normal and leukemic stem cells”, Japan Agency for Medical Research and Development (AMED) [grant number JP24gm6310030] (K.M.), Daiichi Sankyo Foundation of Life Science (K.M.), Mochida Memorial Foundation for Medical and Pharmaceutical Research (K.M.), The Mother and Child Health Found (K.M.), The Shinnihon Foundation of Advanced Medical Treatment Research (K.M.), and the Swedish Research Council (KM). The Lund Stem Cell Center was supported by a Center of Excellence grant in life sciences from the Swedish Foundation for Strategic Research.

