## Supplementary Figures for "Maternal sterol 27-hydroxylase is crucial for securing fetal development"

**Supplementary Figure S1.**

**Supplementary Figure S2.**

**Supplementary Figure S3.**

**Supplementary Figure S4.**

**Supplementary Figure S5.**

**Supplementary Figure S6.**

**Supplementary Figure S7.**

**Supplementary Table 1.**

**Supplementary Table 2.**

**Supplementary Table 3.**

**Supplementary Table 4.**

**Supplementary Movie 1.**

**Supplementary Movie 2.**

**Supplementary Movie 3.**

**Supplementary Movie 4.**

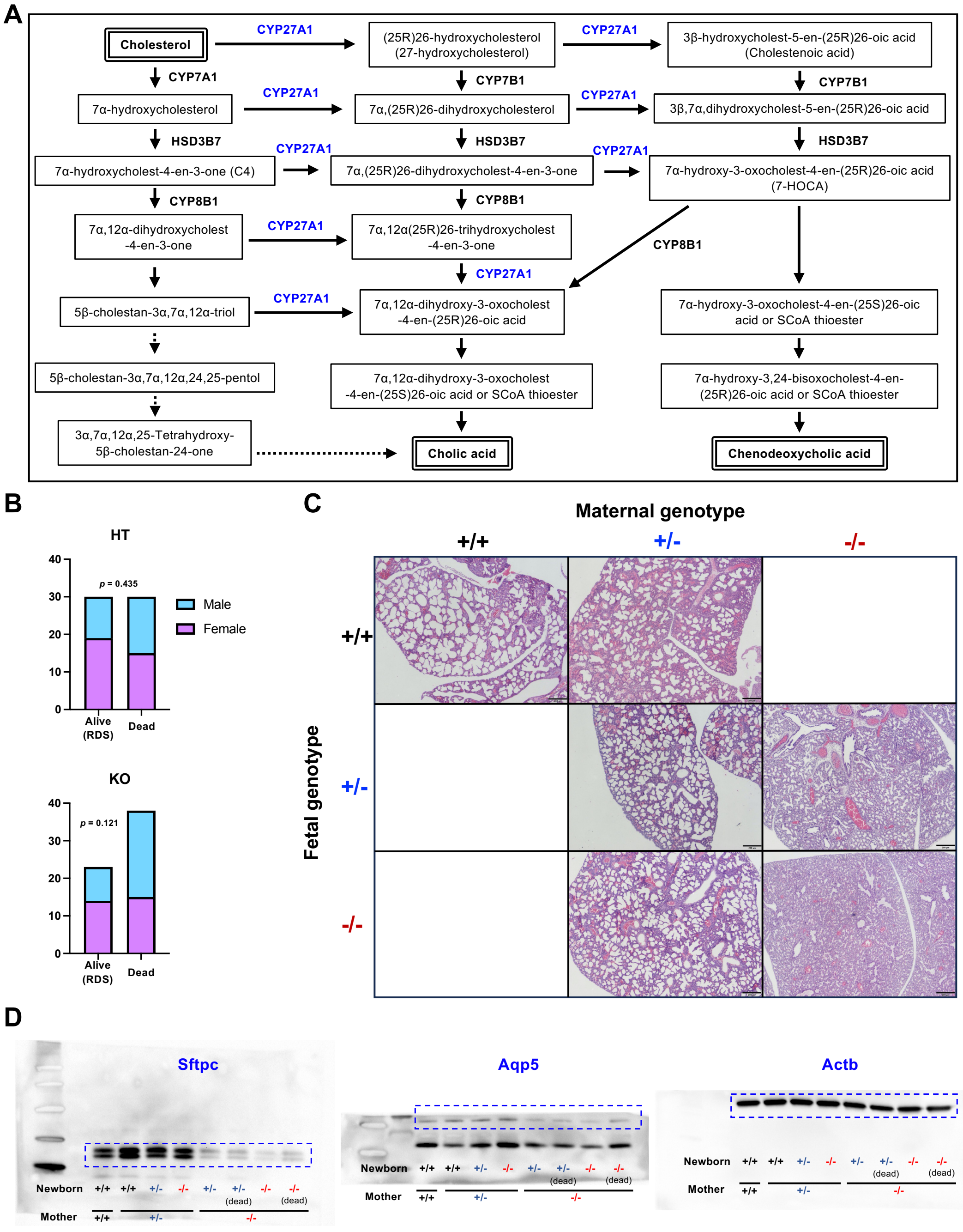


### Supplementary Figure S1.

**(A)** Bile acid synthetic pathways modified from Griffiths et al., 2019. **(B)** Sex ratio in the live births and stillbirths from *Cyp27a1*$^{+/-}$ and *Cyp27a1*$^{-/-}$ mothers. n=4. **(C)** Histology of lungs from newborn mice having various maternal and own genotypes. The scale bar represents 200 μm. **(D)** Western blotting analysis of newborn lungs, corresponding to the Figure 1E.

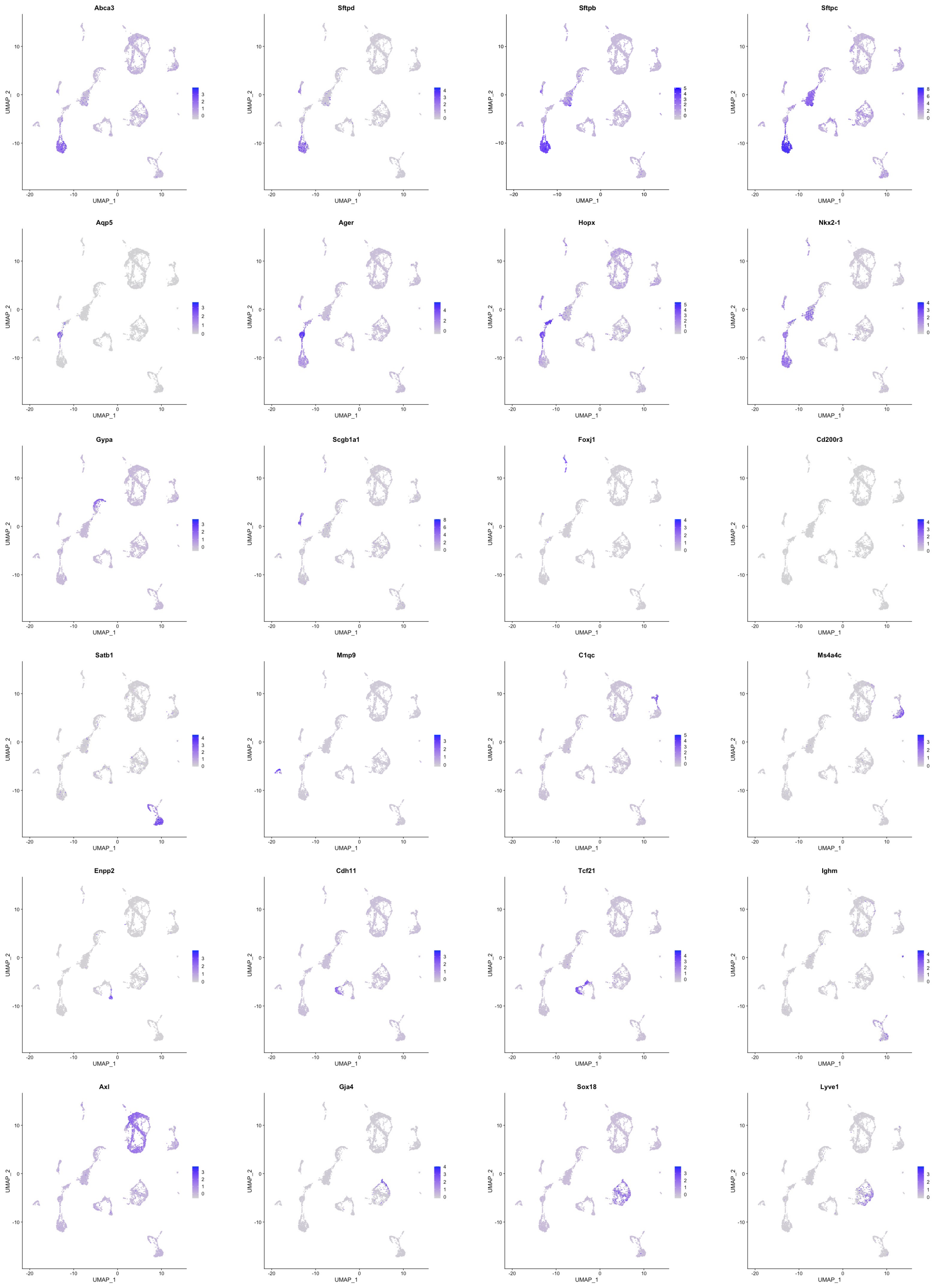


**Supplementary Figure S2.**

Expression patterns of signature genes representing each cluster of E18.5 lungs.

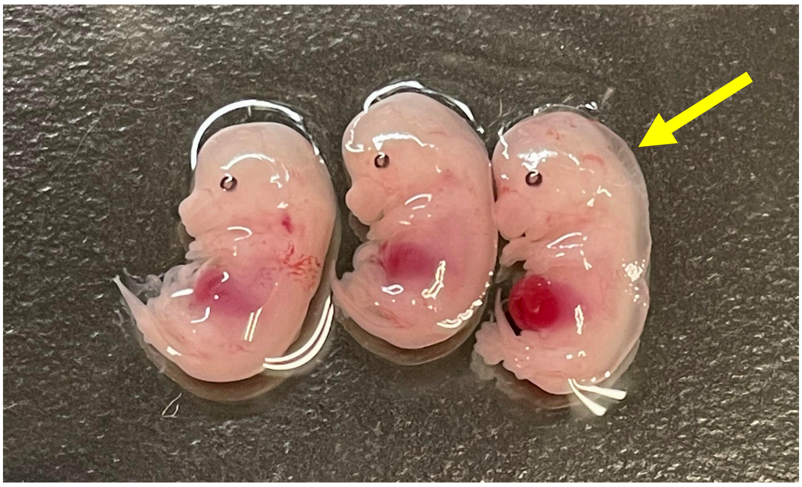

**Supplementary Figure S3.**

A representative photo of E16.5 *Cyp27a1* KO$^{mat}$ fetuses having fetal nuchal edema in their subcutaneous back (indicated by a yellow arrow).

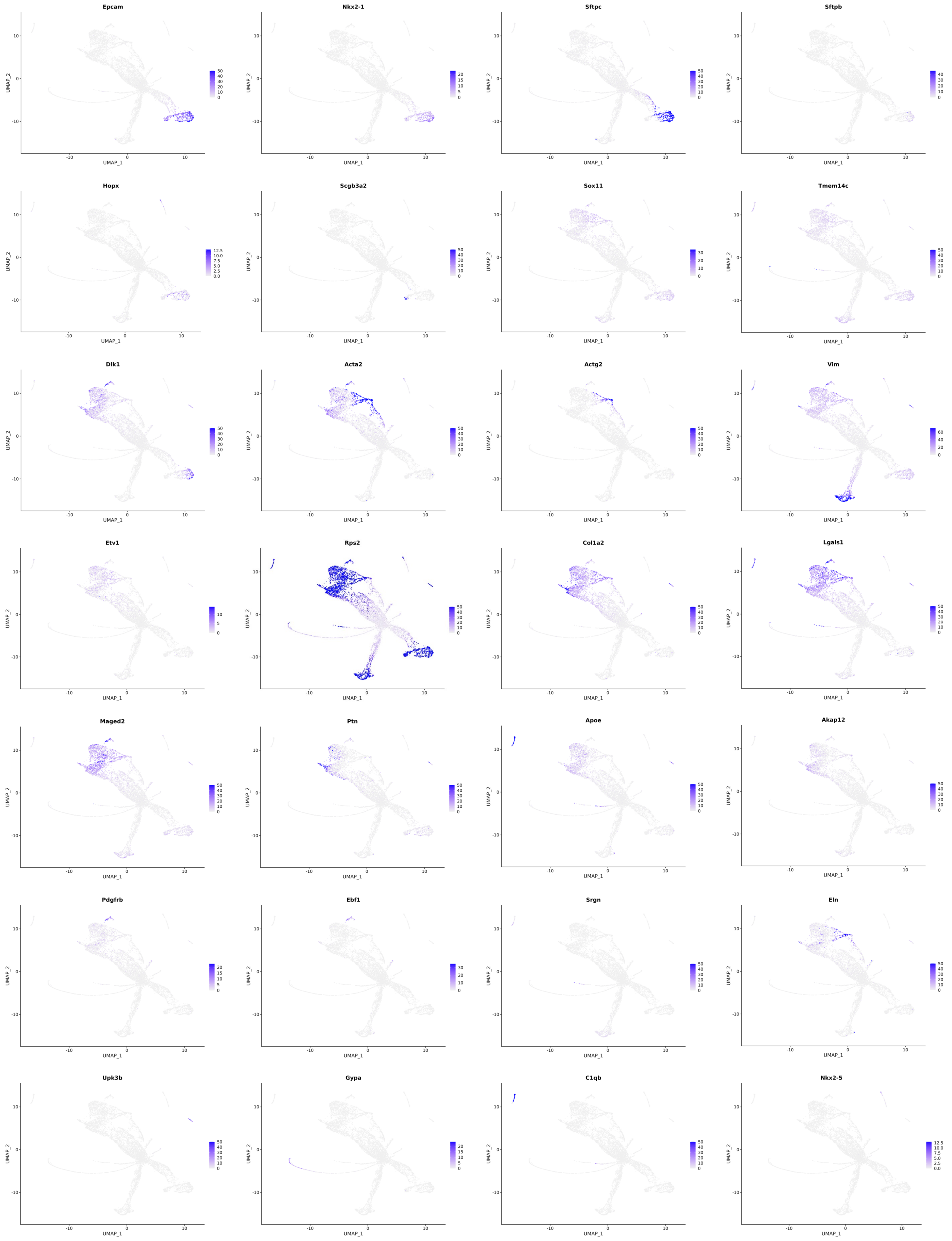


**Supplementary Figure S4.**

Expression patterns of signature genes representing each cluster of E14.5 lungs.

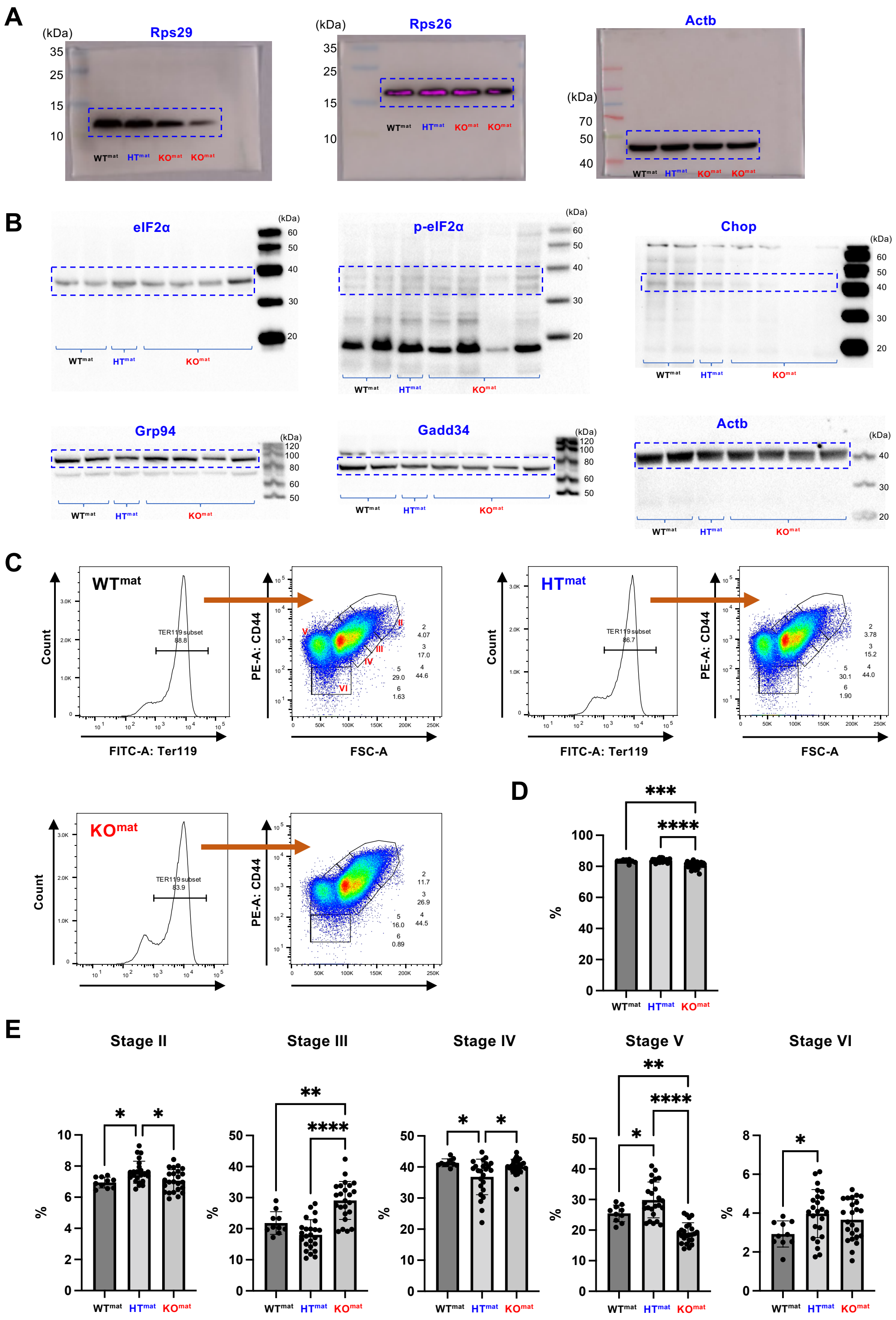


**Supplementary Figure S5.**

**(A)** Western blotting analyses of E16.5 fetal lungs for ribosomal proteinsm corresponding to Figure 3D. **(B)** Western blotting analyses of E14.5 fetal lungs for ER stress markers. **(C)** Representative plots of flow cytometry analysis for the frequencies of different stages of erythroid cells in the liver of E16.5 fetuses. **(D)** The frequency of $Ter119^{+}$ cells. **(E)** Frequencies of each stage of erythroid cells.

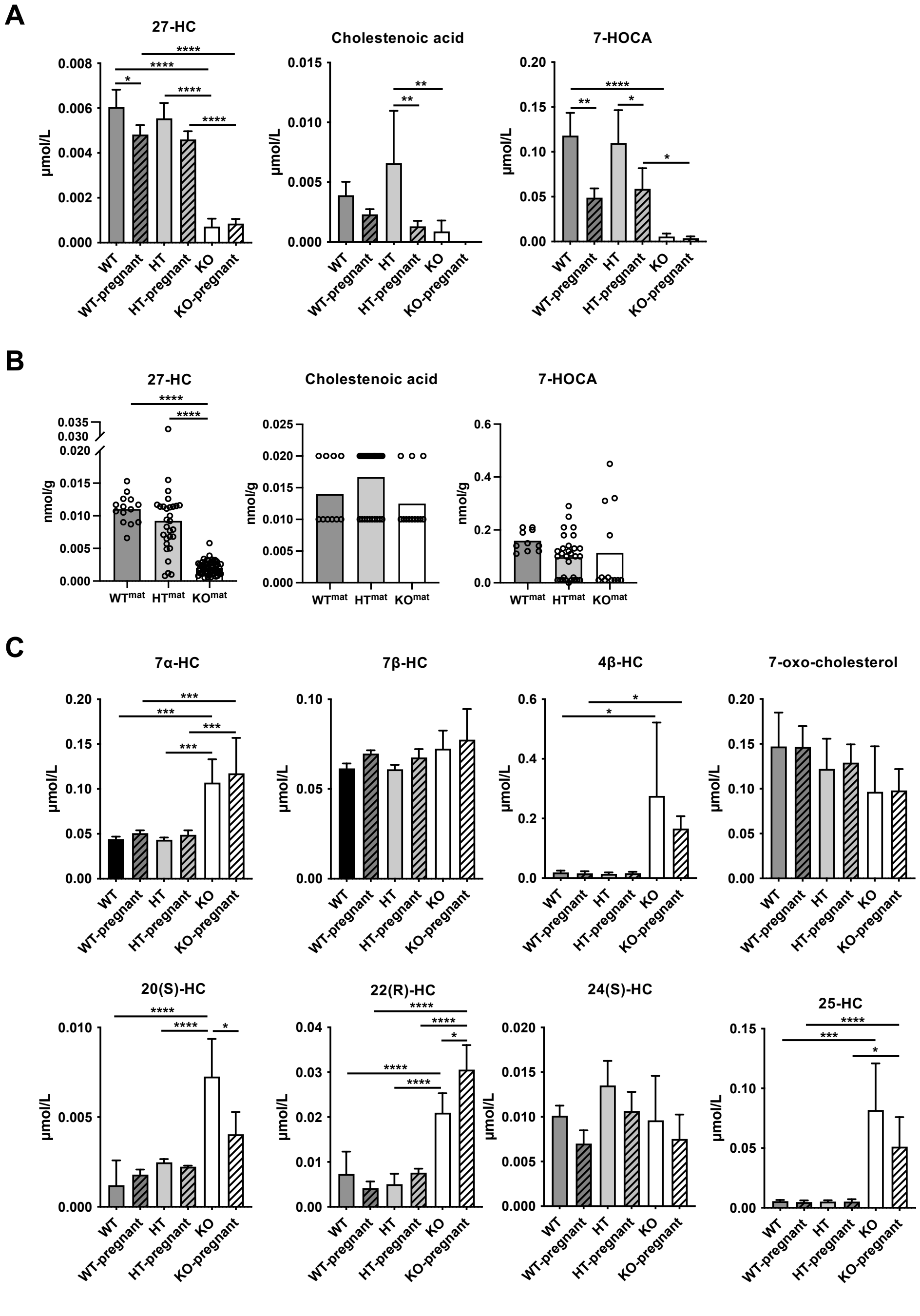


### Supplementary Figure S6.

**(A)** Concentrations of 27-HC and long-chain BAs in the plasms of female mice with or without pregnancy. **(B)** Concentrations of 27-HC and long-chain BAs in the liver of fetuses. **(C)** Concentrations of various oxysterols in the plasms of female mice with or without pregnancy.

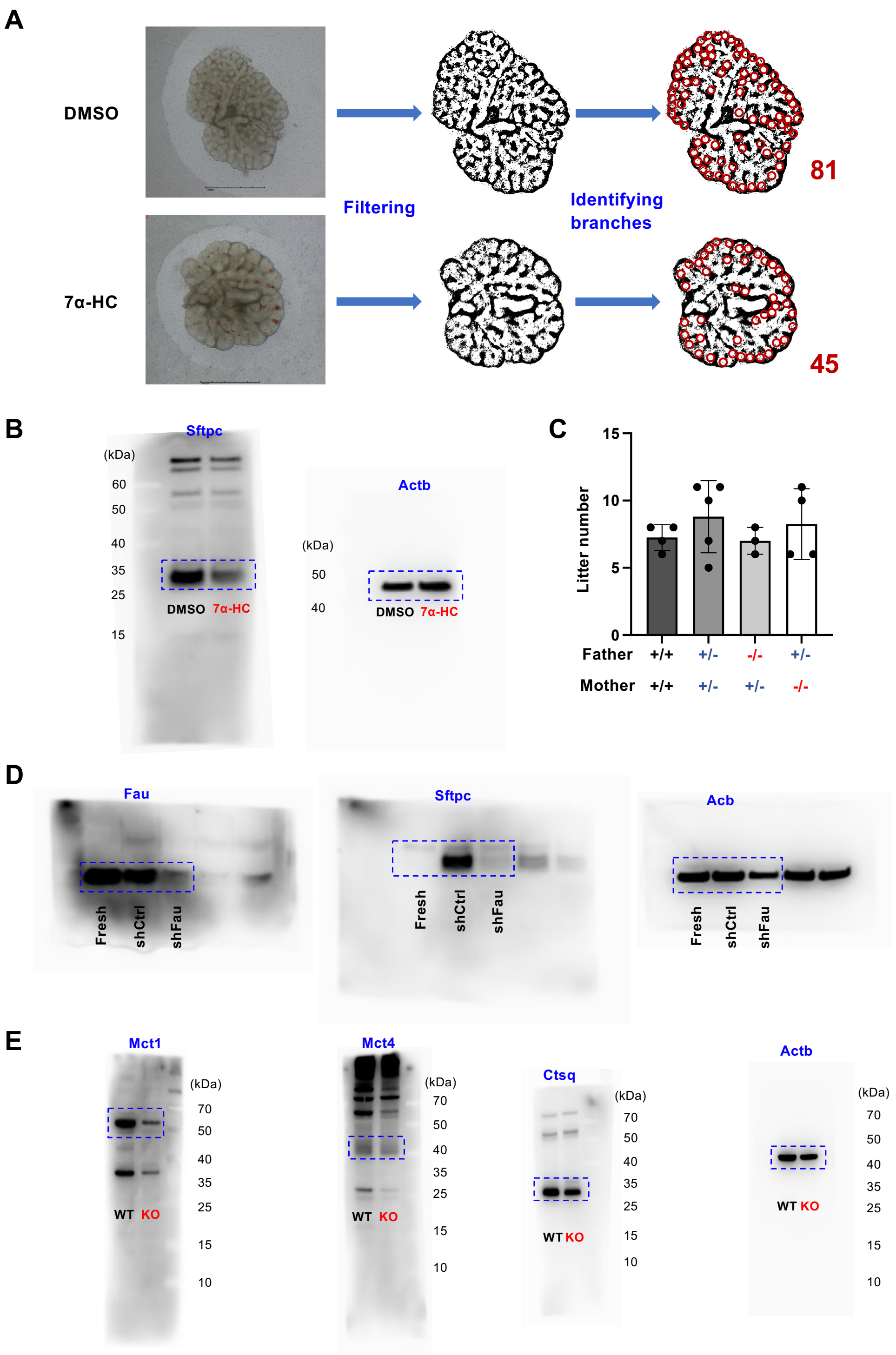


**Supplementary Figure S7.**

**(A)** A schema of alveolar branching counting. The images of *ex vivo* cultured fetal lungs were processed in Photoshop software (Adobe) to extract the branching line using the filtering function. Then the number of branching points were counted. **(B)** Western blotting analyses of *ex vivo* cultured fetal lungs for Sftpc and Actb expression, corresponding to Figure 6G. **(C)** Outcomes from the mating of *Cyp7b1* deficient mice. Wildtype (+/+), heterozygous (+/-), or homozygous (-/-) knockout mice were crossed in various combinations. Litter size was the sum of live births and stillbirths from one pregnant mouse. **(D)** Representative Western blotting analysis of the fetal lungs transduced with lentiviruses, corresponding to Figure 6G. **(E)** Representative western blotting analyses for Mct1 and Mct4 (SynT marker) and Ctsq (sTGC marker), corresponding to Figure 7H.

**Supplementary Table 1.**

A list of signature genes representing each cluster and cluster names of E18.5 scRNA-seq analysis.

**Supplementary Table 2.**

A list of signature genes representing each cluster and cluster names of E14.5 scRNA-seq analysis, and differentially expressed genes in several clusters between $HT^{mat}$ and $KO^{mat}$ fetuses.

**Supplementary Table 3.**

Result of PISA. The quantitative ion intensities (Intensity) were log2-transformed and normalized by the median centering of the values of top 500 precursor in each sample.

**Supplementary Table 4.**

A complete list of abbreviations and names of oxysterols and bile acids analyzed in this study.

**Supplementary Movie 1.**

A representative P1 *Cyp27a1*$^{-/-}$ (KO) newborn mouse delivered from *Cyp27a*$^{+/-}$ (HT) mother.

**Supplementary Movie 2.**

A representative P1 *Cyp27a1*$^{-/-}$ (KO) newborn mouse delivered from *Cyp27a1*$^{-/-}$ (KO) mother.

**Supplementary Movie 3**

*Cyp27a1*$^{-/-}$ (KO) newborn mice generated by IVF using sperm of *Cyp27a1*$^{-/-}$ male mice and egg of *Cyp27a1*$^{-/-}$ female mice.

**Supplementary Movie 4.**

A representative P1 wild-type newborn mouse delivered from wild-type mother injected with 7α-HC.
